# Maternal obesity induces developmental programming of Intestinal stem cells through an IL-17A/PPAR immune-epithelial axis

**DOI:** 10.64898/2026.03.15.711939

**Authors:** Gourab Lahiri, Yesenia Barrera Millan, Swathi Sankar, Karla Mullen, Thomas Hartley McDermott, Dominic R Saiz, Fiona Farnsworth, Matt Torel, Madeline Blatt, Ana Cristina Roginski, Abhigyan Shukla, Esther Florsheim, Benjamin B Bartelle, Khashayarsha Khazaie, Fotini Gounari, Miyeko D Mana

## Abstract

Maternal obesity is associated with increased risk of sporadic colorectal cancer (CRC) in offspring, suggesting that early-life environmental exposures durably shape disease susceptibility. Intestinal stem cells (ISCs), long-lived drivers of epithelial renewal and tumor initiation, are well poised to mediate this effect; however, how maternal obesity influences ISC programming during development remains poorly understood. Using mouse models of diet-induced obesity, we show that exposure to a maternal high-fat Western diet (mHFD) during pre- and postnatal development stably programs colonic ISCs. Offspring exhibit increased ISC proliferation, enhanced self-renewal, a hypermetabolic state, and altered epithelial lineage composition that persists into adulthood despite dietary normalization. These changes are accompanied by increased tumor burden following loss of *Apc* heterozygosity. Mechanistically, we identify the pro-inflammatory cytokine IL-17A as a key extrinsic driver and PPARd/a nuclear receptors as intrinsic mediators of the mHFD phenotype, revealing an immune-epithelial axis that programs ISC function during early life. Together, our findings demonstrate that maternal metabolic environments durably enhance stem cell fitness, providing a mechanistic link between developmental exposure and adult disease risk.

## INTRODUCTION

Early life represents a formative period during which tissue stem cells are specified and mature through maternal-dependent developmental stages. During these windows, environmental exposures can shape molecular patterning and cellular behavior, with lasting consequences for health and disease later in life^1–3^. Despite this importance, the molecular mechanisms underlying developmental programming that establish stem cell fitness remains poorly defined. In adult mice, tissue stem cells can retain memory of prior insults, enabling accelerated responses to subsequent infection or injury challenges^4–9^. Recently, maternal infection was shown to epigenetically remodel fetal intestinal stem cells, resulting in persistent changes in immune-intestinal homeostasis^10^. Further understanding of how maternal-dependent exposures developmentally pattern tissue stem cells will provide critical insights into disease susceptibility and may reveal opportunities to prevent or reverse maladaptive programming.

Intestinal stem cells (ISCs) arise in the endoderm during organogenesis and transition to maturity through introduction of maternal milk and the microbiome^11–13^, compounding the variety of exposures to a developing gut. By weaning, actively cycling *Lgr5*^+^ ISCs promote tissue homeostasis to maintain the epithelia^14^. In adult intestines, ISCs are responsive to signals from the niche and lumen, such that extrinsic cues from diet^15–17^, the microbiome^18^, stroma^19–22^, and immune responses^23,24^ can robustly remodel ISC biology. We and others previously showed that a pro-obesity high-fat Western diet (HFD) promotes stemness and tumorigenicity^17,25^. Mechanistically, a PPAR/CPT1a program mediates these HFD-induced properties, and the PPARs link the obesogenic state to ISC epigenetic reprogramming^9,25^.Yet, these studies of adult exposures overlook the effects in early developmental transformations which we propose could initialize ISCs into pro-tumorigenic conditions *ab initio*.

Maternal obesity is epidemiologically associated with higher CRC risk in adult offspring^26^. The rise of early-onset CRC across multiple countries has focused attention on prenatal and early-life determinants^27,28^. Our previous work supports the hypothesis that *in utero* and perinatal signals may help establish neoplastic trajectories^9,17,25^. Here we investigate whether maternal obesogenic conditions driven by a HFD during maternal-dependent developmental stages could produce long-lasting effects on offspring ISCs. Based on critical developmental processes that occur during ISC fate specification through maturation, we mechanistically test how signals encountered during critical developmental windows durably program ISCs in ways that persist into adulthood.

## RESULTS

### Maternal obesogenic conditions induce persistent effects in offspring ISCs

To assess whether a maternal obesogenic environment imprints long-term alterations in ISCs, we employed a developmental exposure strategy in which dams were mated and maintained on either a control diet or HFD throughout offspring’s gestation and lactation. Offspring were assessed at weaning (d21) or transitioned to a control diet and assessed as young adults (d50) (Figure 1A), a difference of 4 weeks that provides sufficient time for the HFD-induced phenotype to resolve in adults^9,17^. Offspring exposure to a maternal HFD (mHFD) resulted in significantly elevated body weight at both d21 and d50 (Figure 1B), indicative of early metabolic perturbation. To explore the impacts on ISCs, we quantified *Lgr5*^+^ stem cells in the colon using single-molecule RNA *in situ* hybridization (smFISH). mHFD-exposed offspring exhibited a significant increase in *Lgr5*^+^ cells per crypt in both proximal and distal colon at d21, with ISC expansion persisting at d50 (Figure 1C, Figure S1A). The elevation in ISC number was accompanied by increased BrdU incorporation within the crypt base (Figure 1D, Figure S1B), reflecting enhanced proliferative activity. To determine whether these mHFD-induced *Lgr5*^+^ ISCs contribute more robustly to epithelial renewal, we employed lineage tracing using *Lgr5*^CreERT2^; *Rosa26*^TdTomato^ (RFP) reporter mice induced for 40 hr to label ISCs (Figure S1C). Quantification of BrdU^+^RFP^+^ cells, representing actively cycling ISCs, revealed a pronounced increase in mHFD-exposed mice. To test whether ISCs were functionally altered, we conducted *ex vivo* organoid-forming assays on isolated crypts from the proximal and distal colon. At both d21 and d50 timepoints, mHFD-exposed offspring displayed significantly enhanced organoid-forming capacity across colonic regions after 3 days in culture (Figure 1F, Figure S1D). This dual expansion in ISC number and proliferation, coupled with increased ISC regeneration, supports the notion of altered stem cell programming induced by mHFD exposure. To further evaluate the durability of this phenotype, we extended our analysis to d100 mature adults. Overall mass, *Lgr5*^+^ cell abundance, BrdU^+^ proliferating cells, and organoid-forming capacity remained elevated in mHFD-exposed offspring at this later timepoint (Figure S1E,F,G,H), distinct from adult diet-induced obesity which resolves after one month of diet normalization, highlighting that mHFD exposure establishes a stable, self-renewing program during critical windows of fetal and early postnatal development.

**Figure 1.**
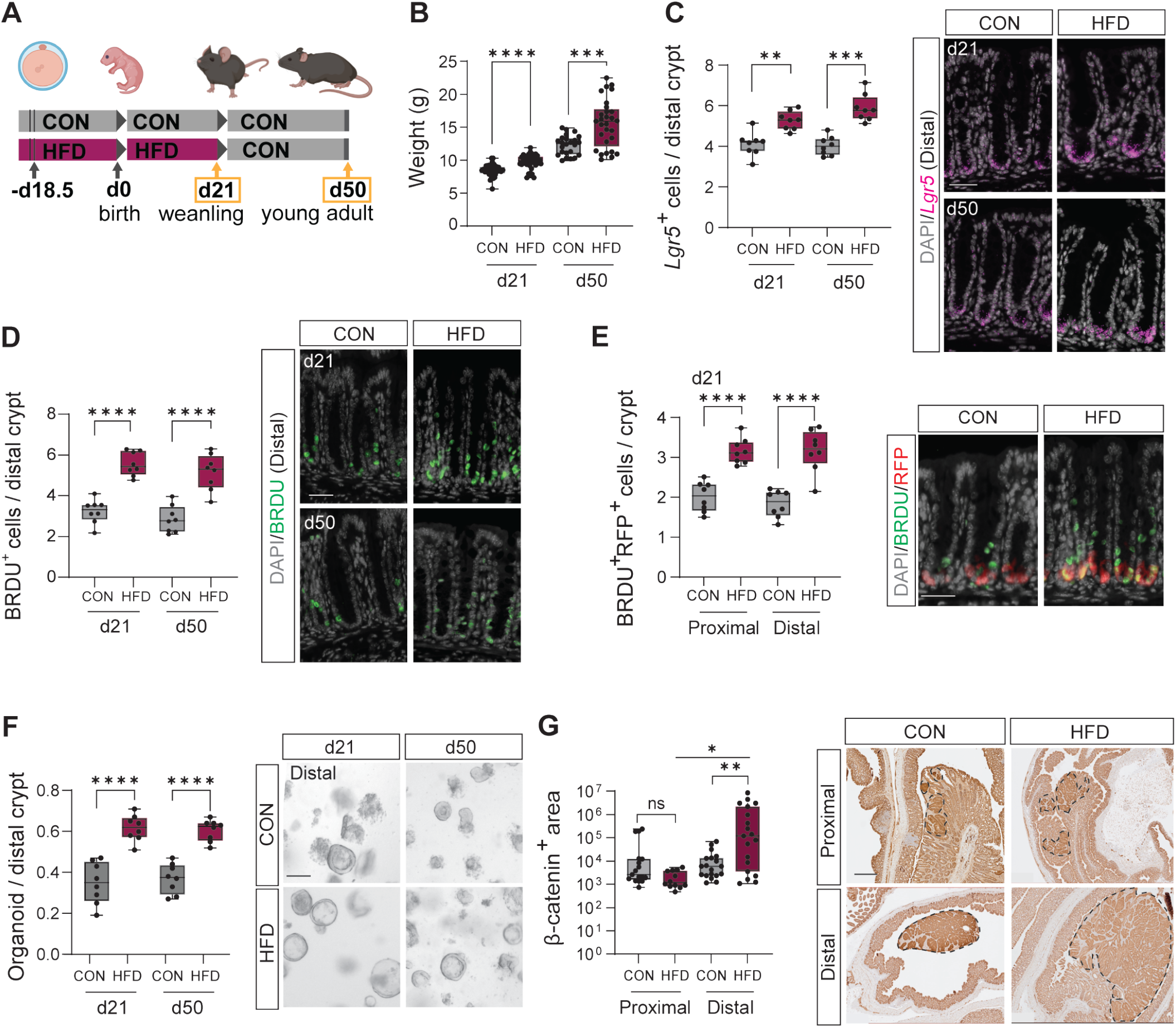
Maternal obesogenic conditions induce persistent effects in offspring ISCs. (A) Experimental schematic depicting the maternal dietary intervention model and analysis timepoints. WT pups born on a maternal high-fat diet (HFD) are switched to the control diet (CON) upon weaning at d21. CON-exposed pups are maintained on the diet until the experimental endpoints. (B) Animal weight at d21 and d50 (n=30). (C,D) Quantification and representative images of *Lgr5*^+^ ISC smFISH (C) and BRDU^+^ cells (D) per crypt from the distal colon at d21 and d50. Each point represents the mean of >20 crypts analyzed per mouse (n=8). Scale bar = 20μm (E) Quantification and representative images of BRDU^+^RFP^+^ proliferative ISCs per crypt from the distal colon at d21. Each point represents the mean of >20 crypts analyzed per mouse (n=8). Scale bar = 20μm (F) Quantification of clonogenic capacity measured by the number of organoids formed per crypt from CON and HFD ISCs. Each point represents the mean of 3+ wells from an individual animal. Scale bar = 10μm. (G) Quantification and representative images of β-catenin^+^ adenomatous lesion area (μm^2^) measured at d120. Each point represents an average lesion area measured from an individual mouse (CON Prox n=17, HFD Prox n=12, CON Dis n=19, HFD Dis n=19). Scale bar = 400μm.

We next asked whether these developmental changes predispose offspring to epithelial transformation. Using an *Apc* loss of heterozygosity model (*Apc*^fl/+^; *Vil*^Cre^), offspring previously exposed to a mHFD during maternal dependence exhibited significantly larger β-catenin^+^ lesions in the distal colon at 4 months (d120) (Figure 1G). Importantly, this greater adenoma area was observed despite the absence of continued HFD exposure, suggesting that the mHFD-programmed ISC state confers elevated oncogenic susceptibility in a regionally biased manner.

### mHFD exposure shifts lineage bias towards a secretory phenotype

To further investigate the persistent changes in ISC properties driven by a mHFD, we performed single-cell RNA-sequencing (scRNA-seq) of distal colonic crypts capturing EpCAM^+^ epithelial and CD45^+^ immune cells across key postnatal timepoints, d21 and d50 (Figure S2A). Quantification and UMAP visualization of epithelial populations at d21 revealed a robust expansion of secretory lineages including goblet cells and enteroendocrine cells (EECs) in HFD-exposed offspring (Figure 2A,B, Figure S2B). The shifts in secretory cell abundance were evident across both developmental stages, indicating that maternal environmental signals impart a persistent lineage adaptation. To validate the transcriptionally inferred secretory bias, we assessed the abundance of functional cell types using immunofluorescence (IF), revealing variable shifts in these populations. Goblet cells (MUC2^+^) are increased across timepoints (Figure 2C, Figure S2C). EECs (PYY^+^) increase at d21 but return to control levels by d50 (Figure S2D,E), reflecting diet-dependent expansion. Deep secretory cells display an inverse pattern in which GFPT1^+^ cells are decreased at d21 but elevated by d50 relative to control counterparts (Figure S2F,G). The overall increase in secretory cell types is supported by the elevated expression level and increased cell frequencies of *Atoh1*, a master transcription factor for secretory cell fate (Figure 2E,F). *Atoh1*^+^ cells per crypt, probed by smFISH, were significantly increased in both colonic regions at d21 and d50 in mHFD-exposed mice (Figure 2D, Figure S2H). *Atoh1* expression level increases in ISCs under mHFD conditions despite less *Atoh1*^+^ ISCs relative to control counterparts (Figure S2I). The abundance of *Atoh1*^+^ cells remained elevated in d100 colons by smFISH, indicating a stabilized commitment to secretory differentiation (Figure S2J). Collectively, these data demonstrate that mHFD-exposure promotes lineage output toward secretory cell fates via durable transcriptional programming.

**Figure 2.**
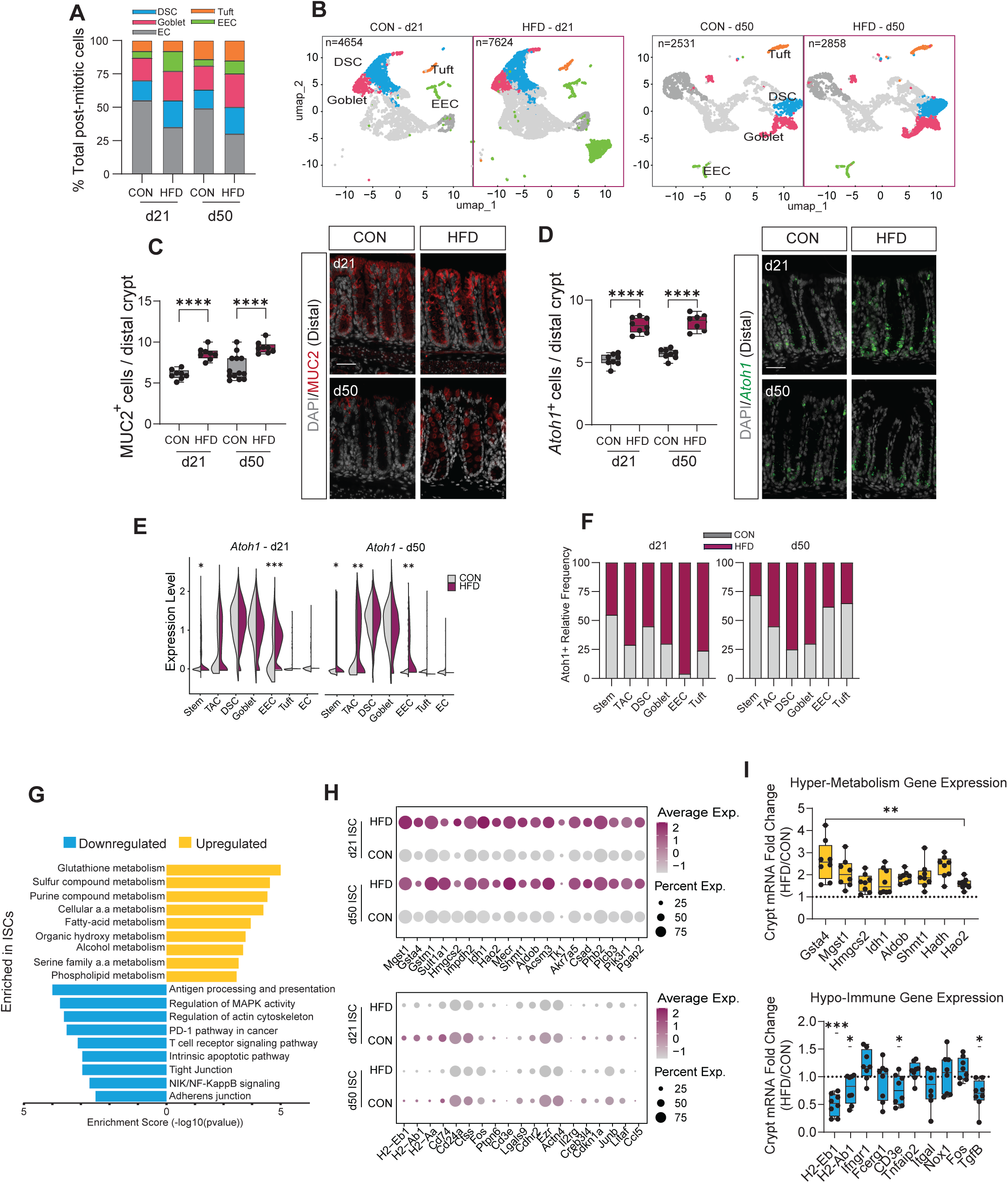
mHFD exposure shifts lineage bias towards a secretory phenotype. (A) Stacked barplot showing the normalized frequency of secretory (DSC, Goblet, EEC, Tuft) and absorptive (EC) cell types across offspring at d21 and d50. (B) UMAP showing the secretory cell clusters in offspring at d21 and d50. (C,D) Quantification and representative images of MUC2^+^ goblet (C) and *Atoh1*^+^ (D) cells per distal colonic crypt at d21 and d50. Each point represents the average mean of >20 crypts analyzed per mouse (n=8). Scale bar = 20μm (E) Violin plot showing the expression levels of *Atoh1* in post-mitotic epithelial cells at d21 and d50. (F) Stacked bar plot showing the relative cell frequency of Atoh1^+^ in CON and HFD at d21 and d50. (G) Pathway enrichment bar plot displaying the top 9 upregulated (sustained increase) and downregulated (sustained decrease) pathways. For DEGs, the log2FC cutoff was set to 0.3 and padj <0.05. (H) Dotplot showing the expression level of genes involved in the Hypermetabolism and Hypoimmune pathways within ISCs from CON and HFD offspring at d21 and d50 (I) qRT-PCR analysis of Hypermetabolism (top) and Hypoimmune (bottom) gene transcription within distal colonic crypts. Each point represents the normalized CT value (HFD/CON) analyzed per d21 mouse (n=8).

To define the stable molecular programs established in ISCs by mHFD exposure, we performed differential gene expression analysis. We confirm the elevated proliferation by promotion of cell-cycle transcripts of S/G2/M markers in mHFD-exposed ISCs (Figure S2K). Next, we focused on genes consistently dysregulated at both d21 and d50, uncovering two temporally sustained groups of transcriptional programs (Figure S2L). Representing stable transcriptional programming beyond early development, we identified a set of genes with sustained upregulation and another set with sustained downregulation (Figure 2H). Pathway enrichment analysis of the sustained upregulated gene set display strong induction of metabolic processes including glutathione metabolism, fatty acid metabolism, and oxidative phosphorylation indicating a shift toward a hypermetabolic epithelial state. Conversely, the sustained downregulated gene set is enriched for antigen-mediated immunity and barrier maintenance pathways, including PD-1 signaling, antigen processing and presentation, tight and adherens junction assembly (Figure 2G), suggesting that HFD-exposed epithelium becomes hyporesponsive to immune signaling and structurally compromised. qRT-PCR analysis of crypts confirmed that multiple genes are upregulated in the hypermetabolic group and downregulated in the hypo-immune set, indicating that stem cell programs contribute to distinct crypt epithelial states (Figure 2I).

### IL-17A cytokine signaling promotes mHFD phenotype

To identify extrinsic factors that may induce these features in a mHFD, we assessed the CD45^+^ immune population. Cytokine gene expression analysis at d21 revealed elevated *Il17a* transcripts across multiple immune cell subsets, with notable expression in γδ T cells (Figure 3A). D50 immune cell counts were too low for analysis. Coincident with increased immune *Il17a*, epithelial expression of *Il17ra* and *Il17rc*, the obligate heterodimers for the IL-17A receptor, were significantly increased broadly in epithelial cells with notable enhancement in ISCs at both d21 and d50, suggesting the formation of a cytokine-responsive epithelium for IL-17A, but not other cytokines (Figure 3B,C, Figure S3A,B). The differential cell abundance expressing *Il17ra* and *Il17rc* is not observed in ISCs from adult 6-month HFD-models (Figure 3D), highlighting a response specific to early-life signals^25^.

**Figure 3.**
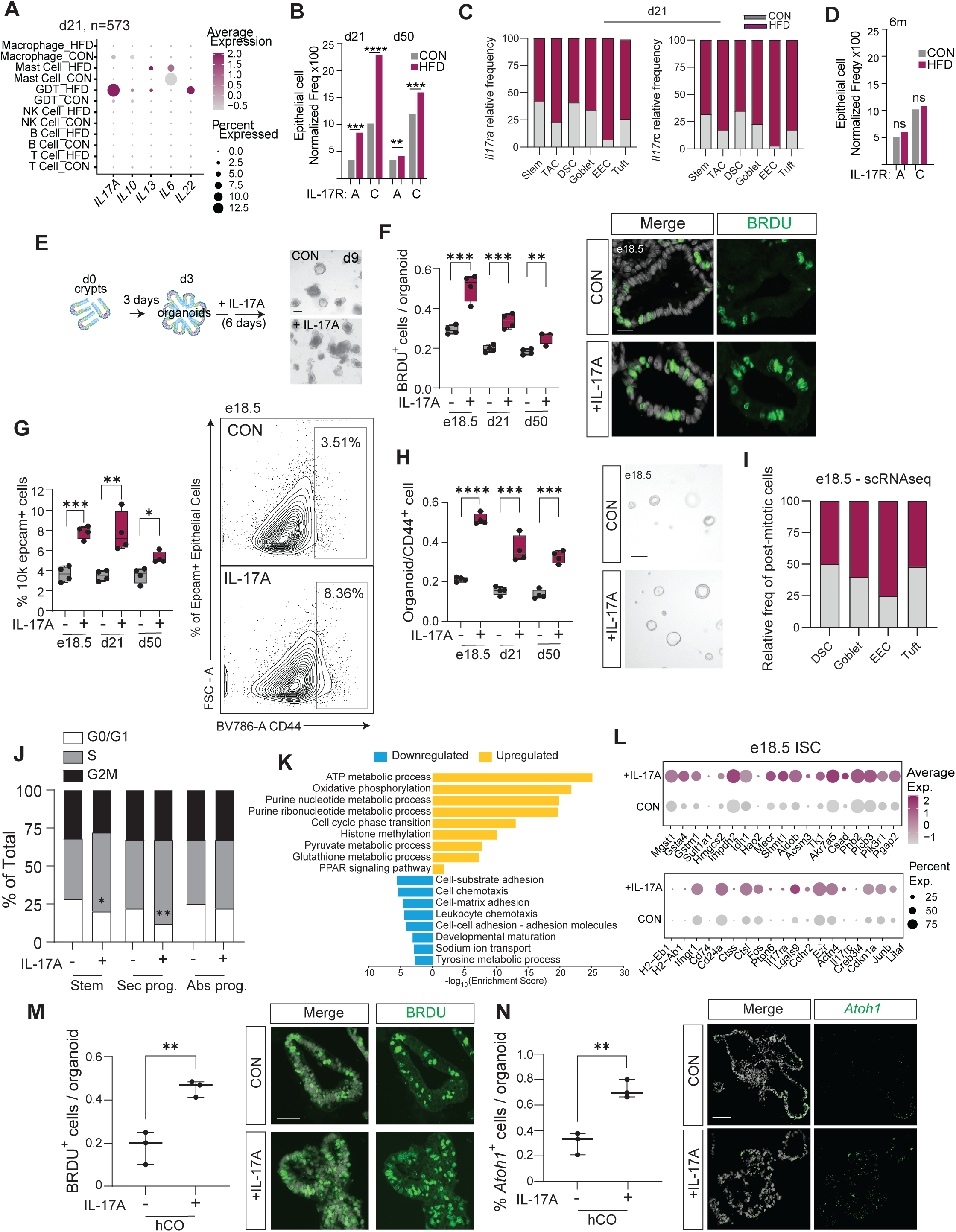
IL-17A cytokine promotes mHFD phenotypes. (A) Dotplot of cytokine expression profiles in the distal immune compartment of offspring at d21 generated from scRNA-seq of 573 cells. (B) Barplot of *Il17ra* and *Il17rc* normalized frequency across total distal colonic epithelium in offspring at d21 and d50 by scRNA-seq of EpCAM^+^ cells. (C) Barplot showing the relative frequency of *Il17ra* and *Il17rc* cells across distal colonic epithelial cell populations in offspring at d21 and d50. (D) Barplot of *Il17ra* and *Il17rc* normalized frequency across total colonic epithelium in adult mice fed a 6 month HFD by scRNA-seq of EpCAM^+^ cells. (E) Experimental schematic depicting *in vitro* IL-17A administration on organoids derived from colonic crypts isolated from WT chow-fed mice at embryonic e18.5, d21, and d50. (F) Quantification and representative images of BRDU^+^ cells per distal colonic organoid (n=4). Each point represents the mean of >10 organoids analyzed per mouse (n=4). (G) Quantification and representative density plot of CD44^+^ cells from Control and IL-17A-treated organoids by flow cytometry across the time points (n=4). (H) Clonogenicity of CD44^+^ sorted cells with representative images, day3 post plating. Each point represents the mean of 3+ wells per mouse. Scale bar = 10μm (I) Stacked bar plot comparing the post-mitotic cell relative frequency from e18.5 IL-17A-treated organoids. (J) Stacked barplot comparing the cell-cycle status of ISCs, Secretory progenitors and Absorptive progenitors in e18.5 IL-17A-treated organoids. (K) Pathway enrichment of top upregulated (sustained increase) and downregulated (sustained decrease) pathways in ISCs from e18.5 IL-17A-treated organoids. For DEGs, the log2FC cutoff was set to 0.3 and padj <0.05. (L) Dotplot of Hypermetabolic and Hypoimmune gene expression in ISCs from e18.5 IL-17A-treated organoids. (M) Quantification and representative images of BRDU^+^ cells per human ESC-derived colonic organoid (n=3). Each point represents the mean of three organoids per differentiation, quantifying BRDU^+^ cells per total DAPI^+^ cells per organoid. Scale bar = 10μm (N) Quantification and representative images of *Atoh1*^+^ transcripts per human ESC-derived colonic organoid. Each point represents the mean of >5 organoids analyzed per hCO differentiation (n=3). Scale bar = 10μm

IL-17A is a pro-inflammatory cytokine that plays a prominent role in promoting protective immunity, but it can also drive inflammatory pathologies, as aberrant IL-17A signaling is associated with chronic inflammation and CRC^29^. To further test if IL-17A is sufficient to induce the mHFD-induced phenotype in ISCs, we administered recombinant IL-17A to cultured colonic organoids derived from e18.5 embryos, thereby testing the effect of the pro-inflammatory cytokine on epithelia preceding direct microbiome influence (Figure 3E). Following 6 days of IL-17A exposure, colonic organoids exhibited a marked increase in BrdU^+^ cells per organoid (Figure 3F), as well as expansion of crypt base EpCAM^+^CD44^+^ cells which enriches for ISCs (Figure 3G), as observed by immunofluorescence and flow-cytometry, respectively. Isolated EpCAM^+^CD44^+^ cells, re-cultured to assess regenerative capacity, robustly formed organoids relative to the control counterpart (Figure 3H). Additionally, IL-17A treatment led to pronounced transcriptional shifts in *Atoh1* and secretory cell type identifiers without increasing the absorptive cell marker *Krt20*, as tested by qRT-PCR (Figure S3C). These mHFD-induced features were also replicated with organoids derived from d21 and d50 pups raised on standard chow, although the induced effects of IL-17A treatment begins to wane as mice mature (Figure 3F-H). Furthermore, recapitulation of the mHFD features appear specific to IL-17A. We tested other cytokines elevated in mHFD immune cells (IL-22, IL-13), known pro-inflammatory effectors (IL-23, Tnfa), and a protective cytokine (IL-10). However, after conducting dose concentration curves (Figure S4A, B), none of these cytokines were able to increase CD44^+^ cells (Figure S4C), enhance organoid forming capacity (Figure S4D) and induce a secretory cell bias (Figure S4E), rendering IL-17A as the candidate cytokine influencing ISCs in offspring exposed to obesogenic conditions.

To assess whether the molecular changes in mHFD ISCs are induced by IL-17A, 6-day treated e18.5 organoids were subject to scRNAseq (Figure S3D). The cell composition of IL-17a-treated organoids confirms changes in lineage allocation with an expansion of goblet cells, EECs, and secretory progenitors (Figure 3I). A greater proportion of ISCs and transient amplifying cells (TACs) appear in S/G2/mitosis phase of the cell-cycle, supporting increased proliferation (Figure 3J). Pathway enrichment analysis of differentially expressed genes in ISCs showed significant upregulation of oxidative phosphorylation, fatty acid metabolism, glutathione metabolism, and cell cycle regulators (Figure 3K). This upregulation was validated by qRT-PCR and consistent with the hypermetabolic phenotype observed *in vivo* (Figure S3E). However, in contrast to the *in vivo* programs, the hypoimmune gene sets in IL-17A organoid treatment were largely elevated or unchanged (Figure 3L, Figure S3F), indicating the inability of IL-17A to downregulate these programs and suggesting other pathways are likely involved to dampen the pro-inflammatory signaling. Overall, IL-17A is largely sufficient to induce key aspects of the mHFD phenotype *in vitro*, providing support that early exposure to pro-inflammatory cues can shape ISC programming (Figure S4F).

To determine whether IL-17a patterning translates to human systems, we directly differentiated human embryonic stem cells to colon organoids (hCOs) with the purpose of mimicking the maternal environment on developing colonic epithelia using an *in vitro* model. We administered IL-17A during mid-to-late hindgut differentiation (d35-42 *in vitro*)^30^ corresponding to the window of mouse intestinal organogenesis and potential maternal IL-17A exposure (Figure S3G,H). Assessment of the mHFD-induced properties showed that IL-17A treatment induced robust proliferation in hCOs seen by increased BrdU incorporation (Figure 3M). Flow-isolated EpCAM^+^CD44^+^ cells from IL-17A treated hCOs had significantly increased organoid formation per 1,000 plated cells (Figure S3I), consistent with enhanced epithelial ISC regeneration. smFISH analysis confirmed increased *Atoh1* transcript levels indicating a shift toward secretory lineage expansion (Figure 3N). This was further supported by qRT-PCR analysis showing increased expression of *Atoh1, Muc2*, and *Chga* (Figure S3J) with no change in absorptive cell markers (*Krt20, Best4*) or mesenchymal marker *Vim*. These findings are consistent with IL-17A as a key early-life effector capable of influencing epithelial stem cell behavior in mammalian species.

### Epithelial *Il17ra* deficiency blocks mHFD phenotype

To test whether IL-17A maintains local or systemic signaling *in vivo*, we measured IL-17A protein levels by ELISA. Serum IL-17A was elevated in mHFD mice at d21 but returned to baseline by d50 after diet normalization, indicating diet-dependent induction systemically (Figure 4A). In contrast, colonic IL-17A levels remained significantly elevated at both stages, particularly in the distal colon (Figure 4B,C), indicating the establishment of a chronic, tissue-localized inflammatory niche. Flow cytometry of mucosal immune cells further demonstrated an expansion of γδ T cells (TCRγδ^+^ RORγT^+^), CD4^+^ T Helper cells, and macrophages (F4/80) at both timepoints in mHFD-exposed colons (Figure S5A,B), confirming that maternal diet leads to the stable expansion of known IL-17A-producing immune cell types^31^. Neutrophils (Ly6G^+^) increase by d50 and NK cells were unchanged, while Treg (Foxp3^+^) and B cell (CD19^+^) abundance decreases. These findings indicate that a mHFD establishes a persistent, IL-17A–enriched inflammatory niche within the colon, sustained by expansion of IL-17A-producing immune populations and shifts in mucosal immune composition.

**Figure 4.**
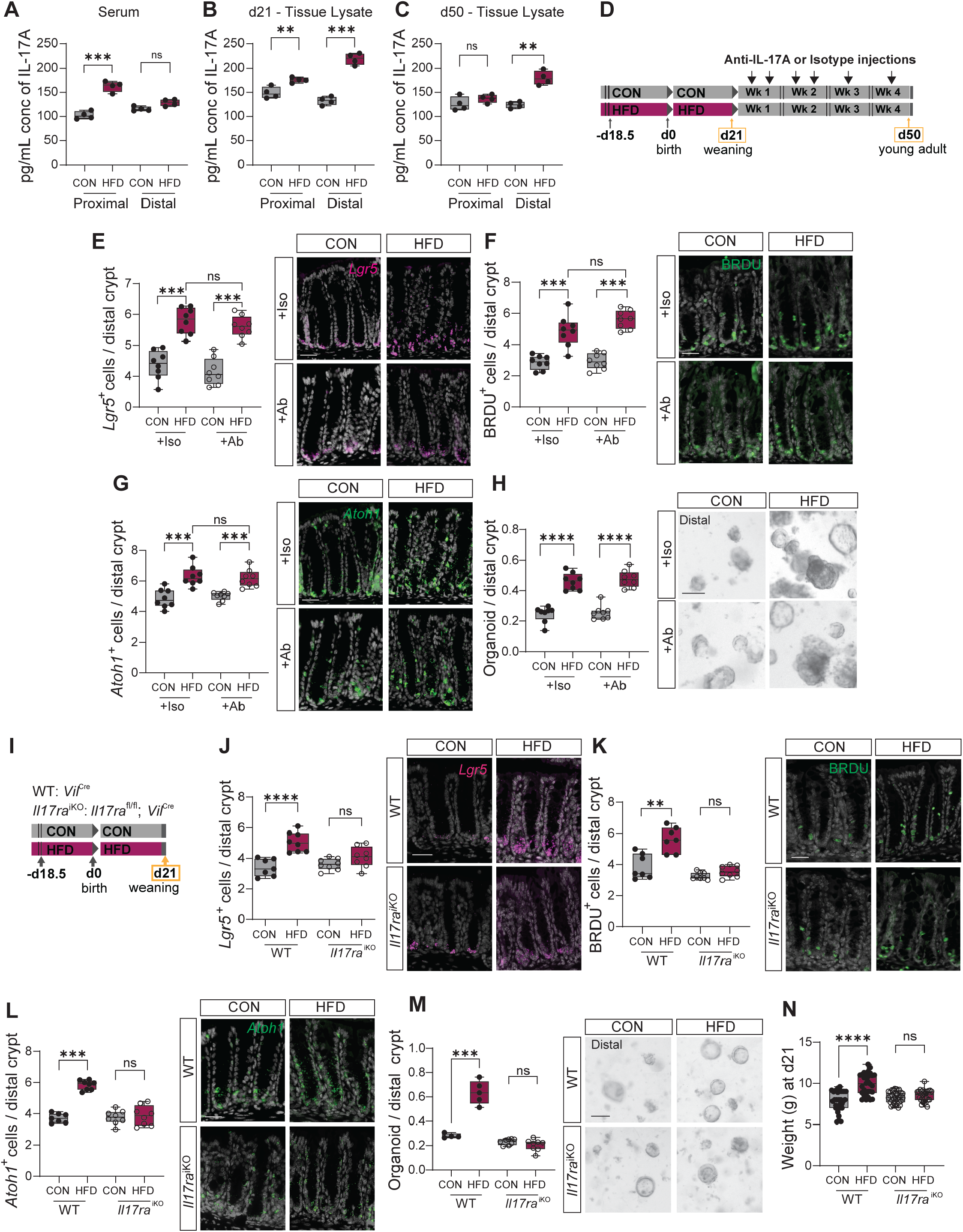
*Il17ra* deficiency blocks mHFD phenotype. (A) Serum IL-17A levels at d21 and d50 via ELISA (n=4). (B,C) Tissue lysate IL-17A levels at d21 (B) and d50 (C) via ELISA (n=4). (D) Experimental schematic depicting the maternal dietary intervention model followed by anti-IL-17A administration post-weaning. Anti-IL-17A or Isotype antibodies were injected IP twice a week for first 2 weeks, followed by once a week for the next 2 weeks. All analyses were conducted at d50 endpoint. (E,F) Quantification and representative images of *Lgr5*^+^ transcripts (E) and BRDU^+^ cells (F) per distal colonic crypt. Each point represents the mean of >20 crypts analyzed per mouse (n=8). Scale bar = 20μm (G) Quantification and representative images of *Atoh1*^+^ transcripts per distal colonic crypt at d50. Each point represents the mean of >20 crypts analyzed per mouse (n=8). Scale bar = 20μm (H) Clonogenicity of distal colonic crypts with representative images. Each point represents the mean of 3+ wells per mouse. Scale bar = 10μm (I) Schematic showing the maternal dietary intervention in d21 WT and *Il17ra*^iKO^ offspring (J) Quantification and representative images of *Lgr5*^+^ ISC transcripts per distal colonic crypt in d21 WT and *Il17ra*^iKO^ offspring. Each point represents the mean of >20 crypts analyzed per mouse (WT CON n=7, WT HFD n=8, *Il17ra*^iKO^ CON n=8, *Il17ra*^iKO^ HFD n=8). Scale bar = 20μm (K) Quantification and representative images of BRDU^+^ proliferative cells per distal colonic crypt. Each point represents the mean of >20 crypts analyzed per mouse (WT CON n=8, WT HFD n=7, *Il17ra*^iKO^ CON n=8, *Il17ra*^iKO^ HFD n=8). Scale bar = 20μm (L) Quantification and representative images of *Atoh1*^+^ transcripts per distal colonic crypt in WT and *Il17ra*^iKO^ offspring. Each point represents the mean of >20 crypts analyzed per mouse (WT CON n=7, WT HFD n=8, *Il17ra*^iKO^ CON n=8, *Il17ra*^iKO^ HFD n=8). Scale bar = 20μm (M) Clonogenicity of distal colonic crypts with representative images from crypt in WT and *Il17ra*^iKO^ offspring from maternal dietary conditions at d21. Each point represents the mean of 3+ wells per mouse. Scale bar = 10μm (N) Weight from WT and *Il-17ra*^iKO^ offspring at d21 (n=30).

One possible explanation for the persistent mHFD-induced phenotype into adulthood is that IL-17A continues to influence the epithelia after diet removal, leading to sustained signaling rather than early molecular ISC programming. To test this hypothesis, we administered anti-IL-17A (Ab) or isotype (IgG/Iso) control antibodies into offspring exposed to a maternal control diet (mCON) or mHFD environment starting at the point of weaning. Animals received injections twice weekly for the first 2 weeks and 1 injection the following 2 weeks until d50 analysis (Figure 4D). ELISA confirmed effective suppression of IL-17A cytokine levels in colonic lysates (Figure S6A,B) and no mass was lost in offspring from either condition due to injections (Figure S6C). However, despite robust cytokine neutralization, the epithelial phenotypes induced by a mHFD exposure were unaffected. Following anti-IL-17A treatment, quantification of *Lgr5*^+^ ISCs (Figure 4E), BrdU^+^ proliferating cells (Figure 4F), *Atoh1*^+^ secretory progenitors (Figure 4G), and MUC2^+^ goblet cells (Figure S6E) showed no significant changes from the mHFD IgG injected controls. Crypt-derived organoid-forming efficiency remained significantly elevated in both the proximal and distal colon in offspring exposed to a mHFD, indicating persistent ISC self-renewal capacity (Figure 4H, Figure S6D) and together, suggesting that post-weaning blockade of IL-17A is insufficient to reverse the functional epithelial programming established during a mHFD. These data imply that the HFD-induced phenotype becomes developmentally fixed under maternal dependent developmental phases and no longer requires continuous IL-17A signaling post weaning.

To test whether IL-17A signaling is required to establish the HFD-induced epithelial phenotype in early life, we used mice harboring an intestinal epithelial-specific knockout of IL-17 receptor A^32^ (*Il17ra*^iKO^: *Il17ra*^fl/fl^; *Vil*^Cre^) (Figure 4I). In this model, epithelial *Il17ra* deletion occurs during fetal colon development, thereby ablating IL-17A responsiveness during the window of mHFD exposure (Figure S6F,G). We observed no changes in colon length or crypt depth (Figure S6H,I). Strikingly, mHFD-exposed offspring lacking epithelial *Il17ra* failed to exhibit any of the hallmark epithelial changes observed in wild-type (WT) mHFD-exposed counterparts. D21 *Il17ra* deficient colons show complete abrogation of the expansion in *Lgr5*^+^ ISCs (Figure 4J), BrdU^+^ proliferative activity (Figure 4K), *Atoh1*^+^and MUC2^+^ secretory cells (Figure 4L, Figure S6K). Organoid-forming capacity remained at baseline mCON levels in both proximal and distal *Il17ra*^iKO^ colonic crypts (Figure 4M, Figure S6J). In addition to the loss of ISC phenotypes, mHFD-exposed *Il17ra*^iKO^ mice did not exhibit the excess body weight gain seen in their WT HFD-exposed counterparts (Figure 4N), suggesting broader metabolic protection arising from disruption of early immune-epithelial signaling. These findings demonstrate that epithelial IL-17RA signaling is essential during a critical developmental window for establishment of the enhanced stemness and secretory program induced by mHFD.

### Intrinsic PPARd/a nuclear receptors are required for the ISC mHFD phenotype

In prior studies, we established the necessity and sufficiency of PPARd/a in mediating adult-onset HFD-induced small intestine and colon ISC phenotypes, including risk of accelerated adenoma formation^9,17,25^. We sought to test the extent to which these nuclear receptors are required for the early-life ISC phenotypes and generated mice possessing intestinal epithelial-specific knockouts of *Ppard* and *Ppara* (*Ppard/a*^iKO^: *Ppard*^fl/fl^; *Ppara*^fl/fl^; *Vil*^Cre^). Loss of these intestinal lipid mediators partially blunts the weight gain in mHFD-exposed offspring (Figure 5A), and we observe no consistent change in colon length or crypt depth in d21 *Ppard/a*^iKO^ offspring (Figure 5B,C). We next screened for the ISC phenotype and bias towards secretory cell fate at d21. In accordance with *Ppard/*a deficiency in adult HFD ISCs, we find that absence of *Ppard/a* in early life blocks the mHFD-induced increase in *Lgr5*^*+*^ ISCs (Figure 5D), BrdU^+^ proliferation, and organoid-forming capacity (Figure 5F,G). Moreover, we find that loss of *Ppard/a* prevents the mHFD-induced increase in MUC2^+^ goblet cells (Figure 5H) and more broadly, *Atoh1*^+^ secretory cells (Figure 5I), indicating an early-life role for these nuclear receptors in regulating lineage composition. Overall, these data identify PPARd/a as critical mediators of mHFD-induced ISC phenotypes in the developing colon.

**Figure 5.**
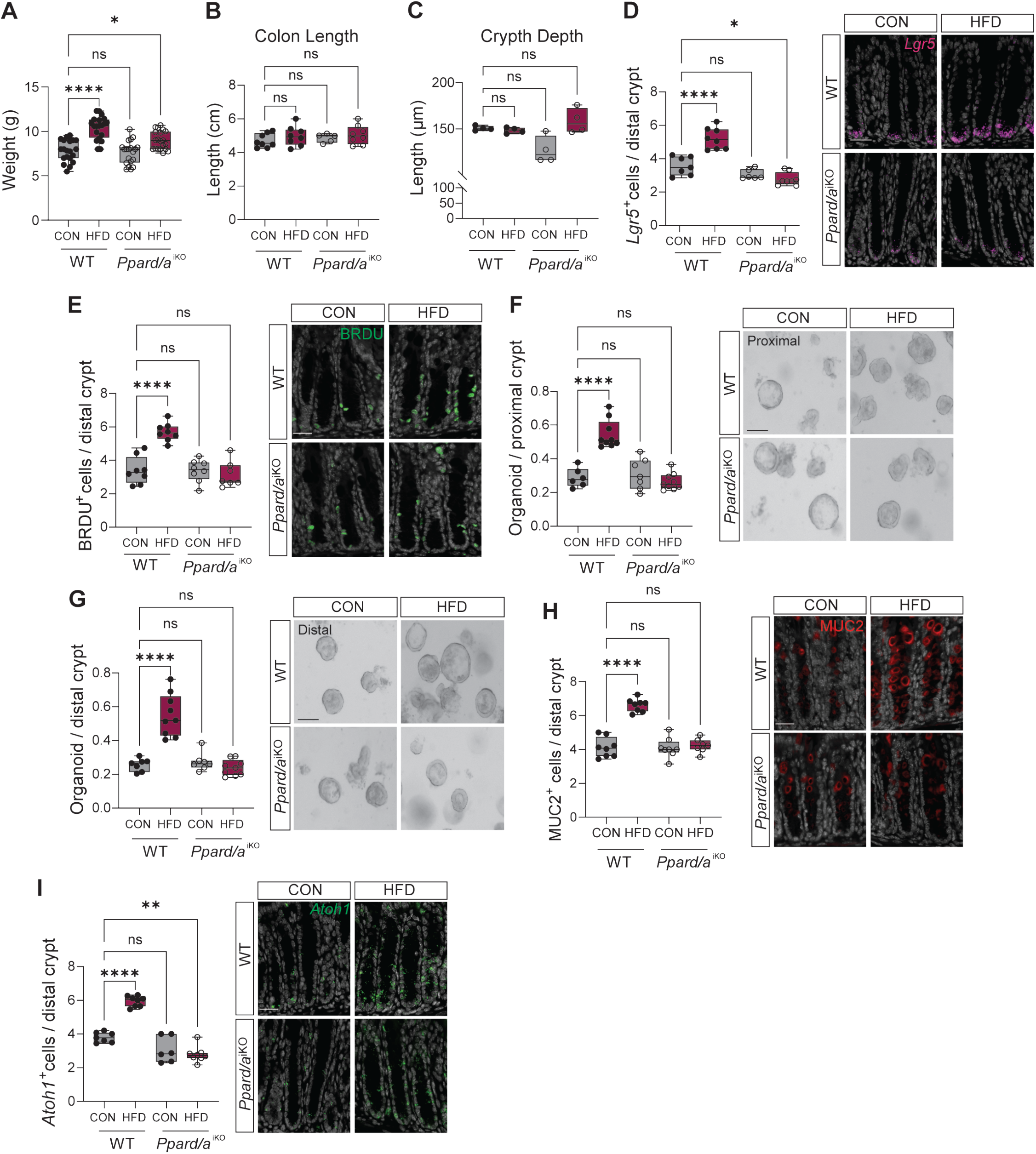
*Ppard/a* deficiency abrogates mHFD phenotype. (A) Weight from WT and *Ppard/a*^iKO^ offspring at d21 (n=20). (B) Quantification of colon length in WT and *Ppard/a*^iKO^ offspring from maternal dietary conditions at d21 (WT CON n=8, WT HFD n=8, *Ppard/a*^iKO^ CON n=5, *Ppard/a*^iKO^ HFD n=8). (C) Quantification of colon crypt depth in WT and *Ppard/a*^iKO^ offspring from maternal dietary conditions at d21 (n=4). (D, E) Quantification and representative images of *Lgr5*^+^ ISC transcripts (D) and BRDU^+^ proliferative cells (F) per distal colonic crypt in d21 WT and *Ppard/a*^iKO^ offspring. Each point represents the mean of >20 crypts analyzed per mouse (*Lgr5*^+^: WT CON n=7, WT HFD n=8, *Ppard/a*^iKO^ CON n=6, *Ppard/a*^iKO^ HFD n=8, BRDU^+^: WT CON n=8, WT HFD n=8, *Ppard/a*^iKO^ CON n=8, *Ppard/a*^iKO^ HFD n=7). Scale bar = 20μm (G) Clonogenicity of proximal colonic crypts with representative images from crypts derived from d21 WT and *Ppard/a*^iKO^ offspring. Each point represents the mean of 3+ wells per mouse. Scale bar = 10μm (H) Clonogenicity of distal colonic crypts with representative images from crypts derived from d21 WT and *Ppard/a*^iKO^ offspring. Each point represents the mean of 3+ wells per mouse. Scale bar = 10μm (I) Quantification and representative images of MUC2^+^ goblet cells per distal colonic crypt in WT and *Ppard/a*^iKO^ offspring. Each point represents the mean of >20 crypts analyzed per mouse (WT CON n=8, WT HFD n=8, *Ppard/a*^iKO^ CON n=8, *Ppard/a*^iKO^ HFD n=7). Scale bar = 20μm (J) Quantification and representative images of *Atoh1*^+^ transcripts per distal colonic crypt in WT and *Ppard/a*^iKO^ offspring. Each point represents the mean of >20 crypts analyzed per mouse (WT CON n=8, WT HFD n=8, *Ppard/a*^iKO^ CON n=4, *Ppard/a*^iKO^ HFD n=4). Scale bar = 20μm

### An IL-17A-PPARd/a axis patterns early ISCs

Given that a mHFD and IL-17A induce a robust metabolic and ISC program, we tested the possibility that the IL-17A response is dependent on a PPAR/CPT1a axis previously identified in adult onset-HFD^25^. We first confirmed that *Ppard* and *Cpt1a* expression is induced upon IL-17A treatment, whereas *Ppara* expression levels appear low and unchanged between conditions, indicating a minimal role in driving the phenotypic or functional outcome (Figure S7A). Next, we used distal colonic organoids derived from WT, *Ppard/a*^iKO^, or *Cpt1a*-deficient mice (*Cpt1a*^iKO^: *Cpt1a*^fl/fl^; *Vil*^Cre^) and treated them with IL-17A cytokine. Loss of *Ppard/a* led to complete abrogation of IL-17A-induced CD44^+^ ISC frequency (Figure 6A), clonogenicity in CD44^+^ sorted cells (Figure 6B), and bias towards the secretory lineage (Figure 6C). *Cpt1a*-deficiency dampened but did not entirely eliminate these features as *Ppard/a*^iKO^ did, indicating that at least some of the IL-17A response functions through more than CPT1a-mediated long-chain fatty acid oxidation. To further substantiate the role of PPARd/a, we assessed the hypermetabolic and hypoimmune programs initially identified in the scRNA-seq and that are robustly upregulated by IL-17A. Here we find that all hypermetabolic genes increased transcriptionally in WT but are blocked in *Ppard/a*^iKO^ organoids (Figure 6D), highlighting its dominant role as a lipid sensor and metabolic effector. Likewise, the hypoimmune genes down-regulated in mHFD offspring and conversely upregulated in IL-17A organoid treatment are also blocked from transcriptional changes in absence of *Ppard/a*, indicating a PPAR-mediated response to IL-17A (Figure 6E). Collectively, the data suggests that the PPARs are required for IL-17A-mediated stemness and induced metabolic activity, and further, may be involved in countering the pro-inflammatory programs *in vivo*.

**Figure 6.**
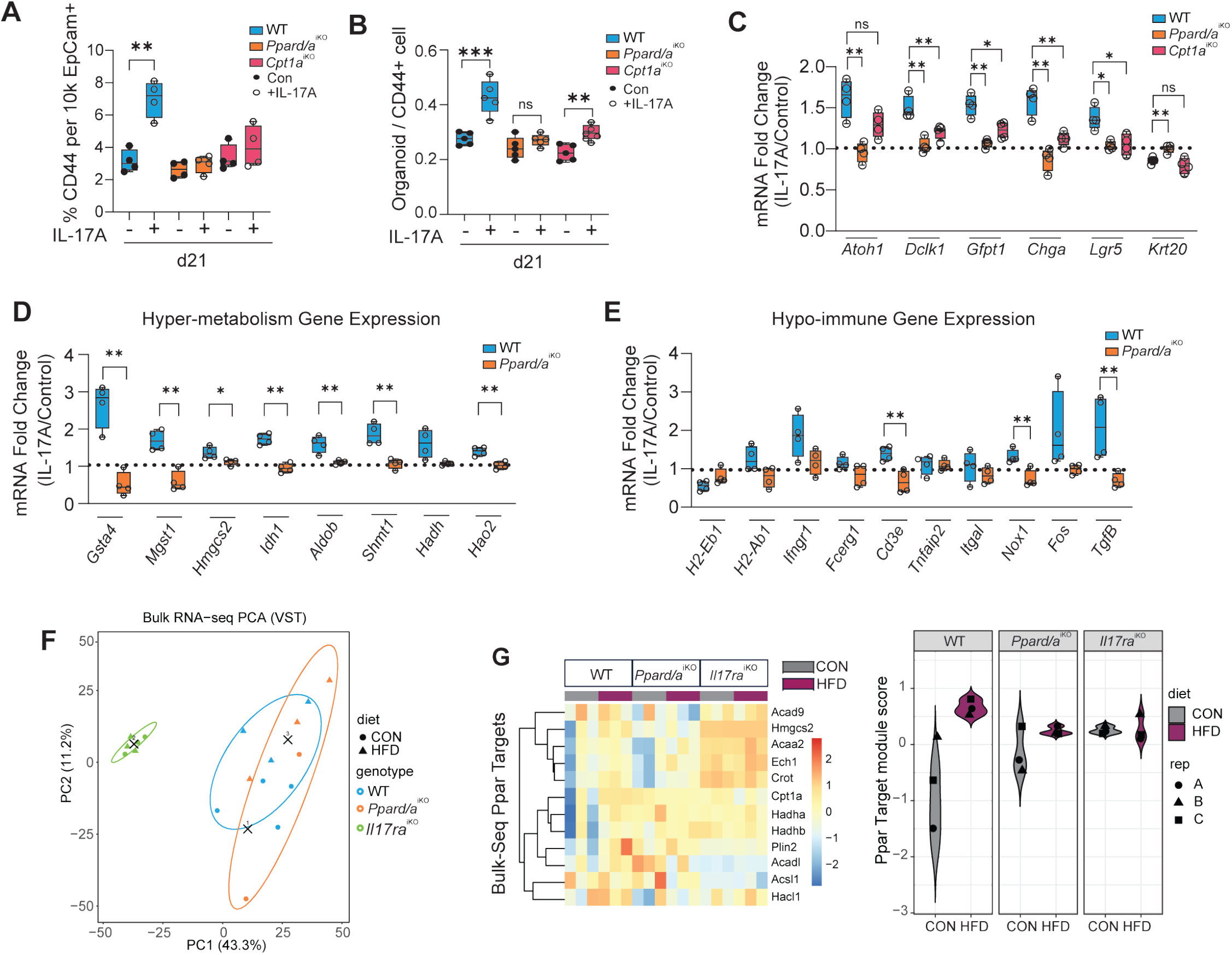
An IL-17A-PPARd/a axis patterns early ISCs. (A) Quantification by flow cytometry of CD44^+^ cells from Control and IL-17A-treated distal colon organoids derived from d21 WT, *Ppard/a*^iKO^ and *Cpt1a*^iKO^ animals (n=4). (B) Clonogenicity of CD44^+^ sorted cells from Control and IL-17A-treated organoids of d21 WT, *Ppard/a*^Iko^ and *Cpt1a*^iKO^ animals. Each point represents the mean of 3+ wells per mouse. (n=4). Scale bar = 10μm (C) qRT-PCR analysis of epithelial cell population from Control and IL-17A-treated mouse distal colon organoids derived from d21 WT, *Ppard/a*^iKO^ and *Cpt1a*^iKO^ animals. Each point represents the normalized CT value (IL-17A/CON) analyzed per mouse (n=4). (D,E) qRT-PCR analysis of Hypermetabolism (D) and Hypoimmune (E) genes from Control and IL-17A-treated mouse distal colon organoids derived from d21 WT and *Ppard/a*^iKO^ animals. Each point represents the normalized CT value (IL-17A/CON) analyzed per mouse (n=4). (F) PCA plot of variance-stabilized gene expression profiles of crypts derived from d21 WT, *Ppard/a*^iKO^ and *Il17ra*^iKO^ offspring. (G) Heatmap analysis showing the z-scored Ppar target gene expression profiles from bulk RNA-Seq of d21 mCON and mHFD crypts derived from WT, *Ppar-d/a*^iKO^ and *Il17ra*^iKO^ mice.

To address whether these relationships translate *in vivo* in offspring exposed to mCON or mHFD conditions, we performed bulk RNA-sequencing on crypts isolated from WT, *Il17ra*^iKO^, and *Ppard/a*^iKO^ mice. Principal component analysis (PCA) separated transcriptional programs into two major groups, with *Il17ra*^iKO^ crypts clustering distinctly from WT and *Ppard/a*^iKO^ samples, suggesting that genotype exerts a stronger influence than diet (Figure 6F). Analysis of the top 500 differentially expressed genes further demonstrated clear separation of *Il17ra*^iKO^ transcriptional programs (Figure S7B), whereas WT and *Ppard/a*^iKO^ crypts showed strong concordance irrespective of dietary exposure. Restricting the analysis to WT and *Ppard/a*^iKO^ crypts revealed a clear genotype-dependent separation, demonstrating that *Ppard/a* independently shapes the transcriptional programs induced *in vivo* (Figure S7C). With further analysis, we find the *in vivo* mHFD-driven hypermetabolic programs exhibit no differential expression between mCON and mHFD within *Il17ra*^iKO^ or *Ppard/a*^iKO^ crypts (Figure S7D). Similarly in the *in vivo* hypoimmune program, no differential response is observed between mCON and mHFD conditions in *Il17ra* deficient crypts (Figure S7E). Given the established role of PPARs in regulating lipid metabolism, we examined expression of canonical PPARd/a target genes. In WT crypts, mHFD induced transcription of PPAR target genes, whereas, this induction was absent in both *Ppard/a*^iKO^ and *Il17ra*^iKO^ crypts, with mHFD samples showing little to no change relative to their control counterparts (Figure 6G). Demonstrating that a PPAR transcriptional program is not induced in *Il17ra* deficient crypts indicates that mHFD-driven activation of PPAR target genes requires active IL-17A signaling. Together we find a cytokine-nuclear receptor axis that links two signaling pathways, typically studied separately, into a single mechanistic pathway that controls the stem cell behavior in early life.

## DISCUSSION

This study explores how a maternal obesogenic environment programs offspring’s ISCs early in life by shaping immune–epithelial interactions through altered molecular circuitry. The irreversible nature of developmental programming resulting in adult-onset diseases that originate from early-life exposure is the core premise of the Developmental Origins of Health and Disease (DOHaD) framework^33,34^. Despite the large body of epidemiological evidence supporting developmental programming, the mechanisms underlying these phenomena remain obscure. Consistent with maternal infection^10^, we find that an early exposure to an obesogenic environment induces durable programmable mechanisms in ISCs. We propose a developmentally restricted IL-17A/PPAR-dependent signaling axis through which a mHFD induces long-lasting changes on offspring’s colonic stem cell behavior and lineage allocation. Our findings establish a mechanistic link between early-life non-pathogenic inflammation and long-term intestinal remodeling, thereby positioning early inflammatory exposures as developmental influencers of durable stem cell states that shape tissue function and disease vulnerability.

Although IL-17A is well known for its role in host defence and inflammation in CRC^35,36^, our data places direct IL-17A signaling as a developmental instructive signal in the gut, capable of influencing ISCs during critical windows of epithelial specification and maturation^37^. We show that fetal ISCs from naive organoids are competent to respond to these immune signals. Whether maternal signals are required during fetal development, post-natal, or both has yet to be clarified^38^. We propose that mHFD-induced inflammatory milieu primes the microenvironment, predisposing tissues to amplified responses upon later inflammatory challenges.

Obesity and high-fat diets are associated with chronic inflammation and altered lipid metabolism. In addition to increased pro-inflammatory IL-17A signaling observed here, we also detect reduced antigen presenting and processing related immune programs, similar to what has been reported in adult-induced HFD models^39^, potentially rendering tumors more evasive of immune surveillance. These findings extend evidence that lipid-rich, PPAR-activated states enhance tumor potential and suggest that early programming of ISCs may render the colon to a persistent pro-tumorigenic environment after exposure to maternal obesogenic conditions. The IL-17A/PPAR axis provides a unifying mechanism, demonstrating how immune and metabolic regulation cooperate to shape tissue stem cells in early life, while highlighting the unresolved question of how PPAR activity stabilizes pro-inflammatory cytokine-induced signaling in ISCs.

Collectively, these findings support a model in which the maternal metabolic state directs immune–epithelial circuitry, establishing a long-lived pro-tumorigenic epithelial landscape. Further investigation of the maternal-offspring dialog will be critical to determine whether this early-life programming can be prevented or therapeutically reversed.

## METHODS

### Experimental Model and Subject Details

Mice were under the husbandry care of the Department of Animal Care and Technologies at Arizona State University (ASU) and the Department of Comparative Medicine at Mayo Clinic Arizona. All procedures were conducted in accordance with the American Association for Accreditation of Laboratory Animal Care and approved by Mayo Clinic’s and ASU’s Institutional Animal Care and Use Committee. The following strains were obtained from the Jackson Laboratory: C57BL/6J (# 000664), IL17RA^fl/fl^ (# 031000), Villin-Cre (Vil^Cre^) (# 004586), Ppard^fl/fl^ (# 005897), Ai9 (Rosa26^TdTomato^. #007909). The following alleles were obtained directly: Ppara^fl/fl^ (Brocker et al., 2017), Lgr5^CreERT2^ (Huch et al., 2013), Cpt1a^fl/fl^ (Schoors et al., 2015), Apc^fl/fl(exon 14)^ (Apc^fl/fl^) (Colnot et al., 2004). The following strains were bred in-house: 1) IL17RA^fl/fl^; Vil^Cre^, 2) Ppard^fl/fl^; Ppara^fl/fl^; Vil^Cre^, 3) Apc^fl/fl^; Vil^Cre^, 4) Lgr5^CreERT2^, Rosa26^TdTomato^, 5) Cpt1a^fl/fl^, Vil^Cre^. Female mice were placed on a high-fat diet (containing 60% kcal from fats (Research Diets D12492i)) 3 days prior to mating and were maintained on the diet throughout gestation and weaning. Control mice were provided a purified control diet (10 kcal% fats, matching sucrose (Research Diets, D12450Ji)). Offspring were analyzed at either d21, d50, or d100. For the d21 cohort, offspring remained on their respective maternal diet until the experimental endpoint. For the d50 and d100 cohorts, all offspring were maintained on Control diet from weaning until the experimental endpoint. Mice used for in vitro organoid assays only were maintained on standard vivarium chow. All offspring mice were sex and age-matched and provided food ad libitum. Mice were kept on a 14/10 day/night light cycle. BrdU (Sigma Aldrich, 19-160) was prepared at 10 mg/mL in PBS (Thomas Scientific, MRGF-6235) and injected at 100 mg/kg 4 hours prior to harvesting tissue. Lineage tracing was done by inducing Cre-mediated recombination by performing an intraperitoneal injection (IP) of tamoxifen (Sigma Aldrich, T5648-1G) suspended in 1 part 200 proof Ethanol (VWR, V1001): 9 parts sunflower seed oil (Spectrum S1929) at a concentration of 10 mg/mL and administered at 100 mg/kg. Antibody neutralization experiments were performed using 10mg/kg concentration of Anti-IL17a (Invitrogen, 16-7173-81) or IgG1 isotype (Invitrogen, MA5-56523) two times a week for the first two weeks post weaning and once a week for the remaining two weeks before the d50 timepoint.

### Crypt Isolation and Culture

Colon was removed, flushed with PBS^−/−^(no calcium, no magnesium), opened laterally, and gently wiped to remove mucus layer. Colon was cut in half to separate proximal and distal sections and incubated at 37°C in PBS^−/−^ + 10 mM EDTA (Invitrogen, 15575020) for 15 min with agitation at 600 rpm. Crypts were then mechanically separated from the connective tissue by shaking and filtered through a 90 μm mesh into a 50 mL conical tube to remove tissue fragments.

For crypt cultures, isolated crypts were counted and embedded in 3.5:6.5 media: Matrigel growth factor reduced (Corning, 356231) mixture at 5-10 crypts per μL and cultured in a crypt culture medium modified from Sato et al. (2009). Intestinal crypts were plated in 10 (5 μl) droplets of Matrigel and placed on flat-bottomed 48-or 98-well plates (Genesee, 25-109 or 25-108). 250 μL (48-well) or 150 μL (96-well) of crypt medium was added to each well and maintained at 37°C in a humidified incubator at 5% CO_2_. Crypt medium was changed every three days and Chir99021 concentration was reduced by half at each passage.

For sorted ISC cultures, following crypt isolation, crypts were resuspended in TrypLE (GIBCO, no phenol red, 12604039) and dissociated into individual cells by heat shocking at 32°C for 90 seconds. Dissociated single cells were stained with the following antibody cocktail for flow cytometry analysis: CD44 (Biolegend, 103041), EpCAM-APC (eBioscience, 17-5791-82), DAPI (BioLegend, 422801). Flow-isolated Epcam^+^ CD44^+^ ISCs were centrifuged at 300 g for 5 min, re-suspended in the appropriate volume of crypt medium and seeded onto 15 μL of Matrigel containing 1 μM JAG-1 protein (Anaspc, AS-61298) in a flat-bottom 96-well plate. 150 μL of crypt medium was added to each well and maintained at 37°C in a humidified incubator at 5% CO_2_.

For in-vitro cytokine treatment, crypts were isolated from the colon and embedded as described above and allowed to form organoids. Three days from plating, the organoids were passaged and treated with the following cytokines separately: Mouse recombinant IL-17a (Thermo, 210-17-25UG), mouse recombinant IL-13 (Thermo, 210-13-10UG), mouse recombinant IL-10 (Thermo, 210-10-10UG), mouse recombinant IL-22 (Thermo, 210-22-10UG), mouse recombinant IL-23 (Thermo, 14-8231-63) and mouse recombinant TNF-a (Thermo, 315-01A-20UG). Dose titration was performed for all the cytokines to find the optimum dosage before treatment on organoids. IL-17a and IL-13 were treated at 5 ng/mL, IL-22 and IL-10 were treated at 10 ng/mL, IL-23 was treated at 7.5 ng/mL and TNFa at 1 ng/mL.

Crypts or flow-isolated CD44^+^ ISCs were grown in Advanced DMEM (GIBCO, 12491015) supplemented with EGF 40 ng/mL (Peprotech, 315-09), conditioned media approximate to 200 ng/mL Noggin and 250 ng/mL R-Spondin, N-acetyl-L-cysteine 1 μM (Sigma Aldrich, A9165), B27 1x (Life Technologies, 17504044), Chir99021 10 μM (LC Laboratories, C-6556), Y-27632 dihydrochloride monohydrate 10 μM (Sigma Aldrich, Y0503). Clonogenicity (colony-forming assay) was assessed on day 3 post plating, unless specified otherwise.

### Immune Cell Isolation and Flow Cytometry

Colon was removed, flushed with PBS^−^/^−^(no calcium, no magnesium), opened laterally, gently wiped to remove mucus layer, cut in half to separate proximal and distal sections, and further cut into 3 mm sections per colon region. Tissue was placed into a 15 mL conical tube containing 10 mL of extraction media composed of 1 mM DTT (Thermo, D1532), 1 mM EDTA, and 10 mL RPMI 1640 Medium (Gibco, 11879020) supplemented with 2% FBS (Gibco, A5670801). Samples were incubated at 37°C for 15 min with agitation at 700 rpm. Sample was filtered through a 40 μm strainer into a 50 mL conical tube and centrifuged at 1500 rpm for 10 min at 4°C. Supernatant was discarded and pellet was resuspended in RPMI 1640 Medium supplemented with 2% FBS. Remaining tissue was resuspended in 10 mL of digestion media containing 1.5 mg Dispase (Sigma Aldrich, D4693), 3.6 mg Type II Collagenase (Thermo Fisher, 17101015), and 10 mL RPMI 1640 Medium supplemented with 2% FBS. Samples were incubated at 37°C for 45 min with agitation at 700 rpm. Samples were filtered into the same 50 mL conical tube using a 100 μm strainer. Samples were centrifuged at 1500 rpm for 10 min at 4°C. Supernatant was removed and samples were resuspended in RPMI 1640 Medium supplemented with 2% FBS. The following antibodies were for the surface marker staining: FITC anti-mouse CD45 (1:500, Biolegend, 103105), BV711 anti-mouse CD3 (1:200, Biolegend, 100241) APC-Cy7 anti-mouse CD4 (1:200, BD, 565650), BV785 anti-mouse CD8a (1:100, Biolegend, 100747), APC anti-mouse CD19 (1:200, BD, 561739), BV650 anti-mouse CD335 (Nkp46) (1:200, Biolegend, 137635), BV650 anti-mouse TCR Gamma Delta (1:200, BD, 564156), PE-Cy5 anti-mouse Ly6G (1:500, Biolegend, 127639), PE anti-mouse CD25 (1:200, Biolegend, 113703), FITC anti-mouse CD45 (1:500, Biolegend, 103107). For intracellular staining we used the Transcription Factor Buffer Set (BD, 562574) and followed the vendor recommended instructions for fixing and permeabilising the cells and the following antibodies were used for the intracellular staining: BV421 Anti-Mouse F4/80 (1:200, BD, 565411), BV711 anti-mouse IL-17A (1:200, Biolegend, 118214) and PE Anti-Mouse ROR-yT (1:200, BD, 562607).

### Histology

All tissues were fixed in 10% formalin (VWR, 77507-022), paraffin-embedded, and sectioned in 4–5-micron sections. Embedding and sectioning were done by the Mayo Clinic Research Histology Core. Antigen retrieval was performed using Borg Decloaker RTU solution (Biocare Medical, BD1000G1) and a pressurized Decloaking Chamber (Biocare Medical, NxGen). Images were acquired using an Olympus VS200 ASW or a Keyence BZ-X800.

#### Immunofluorescence (IF)

The following primary antibodies were used: Rabbit polyclonal anti-MUC2 (1:500, Novus, NBP1-31231), Rat polyclonal Anti-BrdU (1:500, Abcam ab6326), Rabbit polyclonal anti-GFPT1 (1:500, Abcam, ab125069) and Rabbit polyclonal Peptide-YY (1:500, Abcam, ab22663). Nuclear staining was performed using DAPI (1:1000, Biolegend 422801). Alexa Fluor secondary antibodies Anti-Rabbit 568 (1:500, Invitrogen, a10042), Anti-Goat 647 (1:500, Invitrogen, a21447), and Anti-Rat (1:500, Invitrogen, a21208) were used for visualization of all the primary antibodies listed. All primary and secondary antibody dilutions were performed in SignalStain Antibody Diluent (CST, 8112L). Tissue was mounted using Prolong Gold mounting medium (Invitrogen, P36930). Analysis was conducted blind and done using Qupath. >20 crypts per colonic region were analyzed per mouse.

#### Single-molecule fluorescent in situ hybridization (smFISH)

smFISH was performed using Advanced Cell Diagnostics RNAScope™ Multiplex Fluorescent Reagent Kit v2 and according to the manufacturer’s instructions. The smFISH probes used in this study are as follows: Mm-Atoh1-C1 (Ref 408791-C1), Mm-LGR5-C3 (Ref 312171-C3), Hs-ATOH1 (Ref 417861), Hs-LGR5-C2 (Ref 311021-C2). Analysis was conducted blind and done using Qupath. >20 crypts per colonic region were analyzed per mouse. Cells with >5 puncta were considered *Lgr5*^+^ cells. Cells with >10 puncta were considered *Atoh1*^+^ cells.

#### Loss of Heterozygosity

*Apc*^fl/+^; *Vil*^Cre^ offspring were transferred to a Control diet at weaning. Tissue was harvested at d120 post birth. Upon tissue collection, the colon was flushed, and Swiss-rolled prior to fixation. Slides were incubated with 3% H_2_O_2_ for 10 min. A Streptavidin/Biotin Blocking Kit (Vector Laboratories, SP-2002) was used according to manufacturer’s instructions. Slides were stained with β-catenin (1:100, BD Transduction Laboratories, 610154). A Biotin-SP (long spacer) AffiniPure® F(ab’)_2_ Fragment Donkey Anti-Mouse IgG (H+L) secondary (1:500, Jackson ImmunoResearch Laboratories Inc., 715-066-150) was used. Primary and secondary antibody dilutions were performed in SignalStain Antibody Diluent. A VECTASTAIN® ABC-HRP Kit (Vector Laboratories, PK-4000) and SignalStain® DAB Substrate Kit (CST, 8059) was used according to manufacturer’s instructions. Hematoxylin solution Gill I (VWR, 10143-142) and bluing reagent (VWR, 89369-347) counterstains were used. Tissue was mounted using Cytoseal XYL (Electron Microscopy Sciences, 18009). Analysis was conducted using Olympus cellSens software where the adenoma areas were marked by an inbuilt free-hand polyline tool.

### Immunoblotting

Colonic crypts were lysed using TritonX (Sigma Aldrich, X100-500ML) supplemented with 0.5% SDS. Protein was quantified using a Pierce™ BCA Protein Assay Kit and Agilent BioTek Synergy H1 plate reader and according to manufacturer’s instructions. 10 μg of colon crypt lysates were loaded per sample onto a 4%–12% gradient gel (Invitrogen, NP0335BOX), transferred on to PVDF membrane (Immobilon-P transfer, Millipore, IPVH00010), and probed with the following primary antibodies: Rabbit monoclonal anti-Occludin (1:2000, CST, 91131), Rabbit monoclonal Anti-Claudin7 (1:500, Abcam, ab27487), Rabbit polyclonal Anti-Zo2 (1:1000, CST, 2487), Rabbit polyclonal Anti-Claudin3 (1:1000, Thermo, 34-1700). Secondary antibodies were used at a 1:3000 dilution: HRP-linked Anti-Rabbit IgG (CST, 7074S), HRP-linked Anti-Mouse IgG (CST, 7076S). Signal detection was performed using SuperSignal™ West Pico PLUS Chemiluminescent Substrate (Thermo Scientific, 34580) or SuperSignal™ West Femto Maximum Sensitivity Substrate (Thermo Scientific, 34095) and a chemiluminescence (Analytik Jena UVP ChemStudio) system.

### qRT-PCR

Approximately 4,000 crypts were resuspended in 400 μL of TRI Reagent (Sigma Aldrich, 93289). RNA was isolated using a Direct-zol RNA Microprep kit (Zymo Research, R2062) and according to the manufacturer’s instructions. RNA was converted to cDNA, and a qRT-PCR reaction was performed using a Luna Universal One-Step RT-qPCR kit (NEB, E3005) on either of the following systems: QuantStudio™ 3 Real-Time PCR System (96-well), QuantStudio 7 Flex Real-Time PCR System (384-well), or Analytik Jena qTOWERiris (96-well). Primers used are described in the Key Resources table.

### Enzyme-linked immunosorbent assay (ELISA)

Mouse serum and colon tissue lysates were analyzed for IL-17A protein levels using the Mouse IL-17A ELISA Kit (Proteintech, KE10020) according to the manufacturer’s instructions. Serum was obtained by cardiac puncture and centrifuged at 10,000 g for 30 min at 4°C. Supernatants were aliquoted and stored at ^−^80°C until use. For tissue measurements, distal colon segments were rinsed in ice-cold PBS and homogenized in RIPA Lysis and Extraction Buffer (eBiosciences, 89901) supplemented with Halt™ Protease Inhibitor Cocktail (100X) (Thermo Scientific, 78430) on ice. Homogenates were clarified by centrifugation at 10,000 g for 30 min at 4°C, and supernatants were collected. Protein concentration was measured by BCA assay, and lysates were normalized to 1 mg/mL total protein prior to ELISA. Samples and standards were run in duplicate. Standard curves were generated using recombinant IL-17A provided with the kit. Serum was assayed at 1:2–1:5 dilutions, and tissue lysates at 1:2–1:10 dilutions, to ensure measurements fell within the assay’s dynamic range. Raw absorbance values were background-corrected and fit to a four-parameter logistic regression curve (4-PL) using GraphPad Prism. Concentrations were interpolated from the standard curve and adjusted for dilution factors. Final values were expressed as pg/mL for serum or pg/mg total protein for tissue lysates.

### Human Embryonic Stem Cell (hESC) Culture and Differentiation to Colon Organoids

hESC maintenance. H1 human embryonic stem cells (hESCs) were maintained on 6-well plates (Genesee, 25-105) coated with hESC-qualified Matrigel (Corning, 354277) in mTeSR Plus medium (StemCell Technologies, 100-0276). Media was changed every other day. For passaging, colonies were dissociated using ReLeSR (StemCell Technologies, 100-0483) and re-plated at a 1:6 ratio.

Directed differentiation. Differentiation toward human colon organoids (hCOs) was performed using the STEMdiff Intestinal Organoid Kit (StemCell Technologies, 05140). hESCs were seeded as small clumps onto 24-well plates (Genesee, 25-107) coated with hESC-qualified Matrigel. At 24–48 hours post-seeding, cells were cultured in warmed Endodermal Basal Medium (StemCell Technologies, 05111) supplemented with CJ (StemCell Technologies, 05113) to induce definitive endoderm for three days. Media was refreshed daily. Cells were then cultured for six additional days in Endodermal Basal Medium supplemented with PK (StemCell Technologies, 05141) and UB (StemCell Technologies, 05142) to drive hindgut specification. Floating spheroids were collected on day 7–9, filtered, and transferred into Matrigel domes in 24-well plates.

Colon organoid generation and maintenance. For hCO induction, spheroids were cultured in hCO medium (Table 1) supplemented with 95 ng/mL BMP2 for three days, after which BMP2 was withdrawn. For human intestinal organoids (hIOs), spheroids were maintained in Intestinal Organoid Basal Medium (StemCell Technologies, 05143) supplemented with Intestinal Organoid Supplement (StemCell Technologies, 05144) and L-glutamine. Media was changed every three days, and organoids were passaged every seven days.

Organoid passaging. All tubes in contact with organoids were pre-coated with Anti-Adherence Solution (StemCell Technologies, 07010). Organoids were washed with DPBS (GIBCO, 14190-144), collected in DMEM/F12 (Thermo Fisher, 11330-032), and allowed to settle on ice by gravity. After removal of single-cell debris, organoids were centrifuged at 300 g for 5 min, resuspended in 50 µL Matrigel, and plated into 24-well plates with complete hCO medium.

IL-17A treatment. To assess IL-17A effects, hCOs were treated with recombinant human IL-17A (Peprotech, 200-17-5UG). A concentration curve (5, 10, 25, 50, and 100 ng/mL) was initially tested. Based on RT-qPCR readout for the secretory lineage transcription factor Atoh1 and absence of cytotoxicity, 10 ng/mL IL-17A was selected for experiments. IL-17A was added to hCO media starting on day 25. Organoids were maintained with media changes every three days and passaged weekly and collected on d42 for downstream analyses.

Collection for histological analysis. BrdU was prepared at 10 mg/mL in PBS. Organoids were pulsed with BrdU for 1 hour prior to being collected, fixed with 4% paraformaldehyde, paraffin-embedded, and sectioned in 4–5-micron sections. Embedding and sectioning was done by the Mayo Clinic Research Histology Core.

### scRNA-seq data processing and differential expression analysis

Single-cell RNA sequencing data were acquired for Epcam+ Epithelial and CD45+ Immune cells isolated from the distal colon of control and high-fat diet mice. The samples were processed using a 10x Genomics Chromium Single Cell Controller. To produce paired-end reads, the resulting libraries were sequenced on an Illumina sequencing platform. Using Cell Ranger (10x Genomics), the raw sequencing data were initially processed for transcript alignment, barcode processing, and unique molecular identifier (UMI) counting. After that, R was used to import the filtered feature-barcode matrices for the high-fat and control diet samples using the “Read10X” function from the “Seurat” package. To do initial quality control, Seurat objects were created for each condition, and cells were filtered using the following criteria: fewer than 60,000 total UMI counts to eliminate dead cells or doublets, and cells expressing at least 200 genes. Additionally, the number of mitochondria was measured, and cells containing more than 15% mitochondrial genes were not included in the analysis that followed. Post Normalization and integration, principal component analysis (PCA) was used to minimize dimensionality, keeping the first 50 principal components after scaling the data. The data was then shown in two dimensions by building a UMAP (Uniform Manifold Approximation and Projection). Using a graph-based clustering technique, clusters were found, and the granularity of the clustering was adjusted by setting the resolution parameter to 0.5. To find marker genes for each cluster in which at least 25% of the cells in either of the examined clusters expressed the gene, differential expression analysis was performed. The identification of unique cellular identities and states resulting from the food intervention was made easier by this approach. Based on information from the literature and the expression of known marker genes, cell clusters were identified.

### Bulk RNA-seq library preparation and sequencing

Approximately 4,000 crypts were resuspended in 400 μL of TRI Reagent. RNA was isolated using a Direct-zol RNA Microprep kit (Zymo Research, R2062) and according to the manufacturer’s instructions. Library prep and sequencing was conducted by Novogene using an Illumina NovaSeq platform to generate 150-bp paired-end reads (PE150).

### Bulk RNA-seq data processing and analysis

Bulk Raw sequencing reads were assessed for quality using FastQC, and adapter sequences and low-quality bases were trimmed using Trim Galore with default parameters. Cleaned reads were aligned to the mouse reference genome (GRCm39) using HISAT2 (v2.0.5) with default settings and gene annotation from GENCODE (release MXX). Only uniquely mapped reads were retained for downstream analyses. Gene-level read counts were generated using FeatureCounts (v1.5.0-p3) was used to count the reads numbers mapped to each gene. Then FPKM of each gene was calculated based on the length of the gene and reads count mapped to th<u>e</u> gene. This count data was imported into R and processed using DESeq2. Raw gene-level counts were cleaned by removing annotation columns, converting values to numeric integers, and standardizing ENSEMBL gene identifiers by trimming version suffixes. Differential expression analysis was performed using DESeq2 with a global design formula to model main effects and interactions. Variance stabilizing transformation (VST) was applied for downstream visualization. Marker-focused heatmaps were generated using pheatmap from VST-normalized expression values, scaled by row, and annotated by genotype, diet, and replicate.

### Quantification and statistical analysis

Unless otherwise specified in the figure legends, all experiments reported in this study were repeated at least five times. Unless otherwise specified in the main text or figure legends, all sample numbers (n) represent biological replicates. For murine organoid assays, 2-5 wells per mouse per ex vivo treatment were analysed. All center values shown in graphs refer to the mean. Sample size estimates were not used. Animals were randomly assigned to groups. Experiments used roughly equivalent male and female mice to avoid sex bias. Studies were not conducted blind, with the exception of all histological analyses. Please note that statistical details are found in the figure legends.

### Software

Histological and immunofluorescence images were acquired using Olympus VS200 ASW or a Keyence BZ-X800. Image processing and quantitative analyses, including cell counting, area measurements, and fluorescence intensity quantification, were performed using Olympus cellSens software, QuPath and ImageJ (Fiji). Flow cytometry data were analyzed using FlowJo for gating, population identification, and quantification of marker expression. Statistical analyses and graphical representations were generated using GraphPad Prism. Figures and schematic illustrations were prepared using BioRender.

## RESOURCE AVAILABILITY

### Lead contact

Further information and requests for resources and reagents should be directed to the lead contact, Miyeko D Mana

## ACKNOWLEDGEMENTS

Cell sorting was assisted by Adam Kindelin and the ASU Biosciences Flow Cytometry Core using BD FACSymphony S6, acquired by the NIH SIG award 1-S10-OD032287-01. ASU Research Computing provided HPC and storage support ^40^. BBB is supported by NIBIB-R21EB034970, NIMH-DP2MH136493. MDM is supported by NCI-K22CA241083, RSG-22-095-01-CCB, NCI-R01CA301086, NIDDK-R01DK143944, RFGA2024-022-020.

## AUTHOR CONTRIBUTIONS

Conceptualization: GL, KM, MDM; Methodology: GL, BBB, MDM; Formal Analysis and Investigation: GL, YBM, SS, KM, THM, DRS, FF, MT, MB, AS, ACR; Writing-original draft: GL; Supervision: EF, BBB, KK, FG, MDM; Writing-Review and editing: All authors; Funding acquisition: MDM

