## Supplementary figures and images for "Maternal obesity induces developmental programming of Intestinal stem cells through an IL-17A/PPAR immune-epithelial axis"

### Supplementary Figure 1

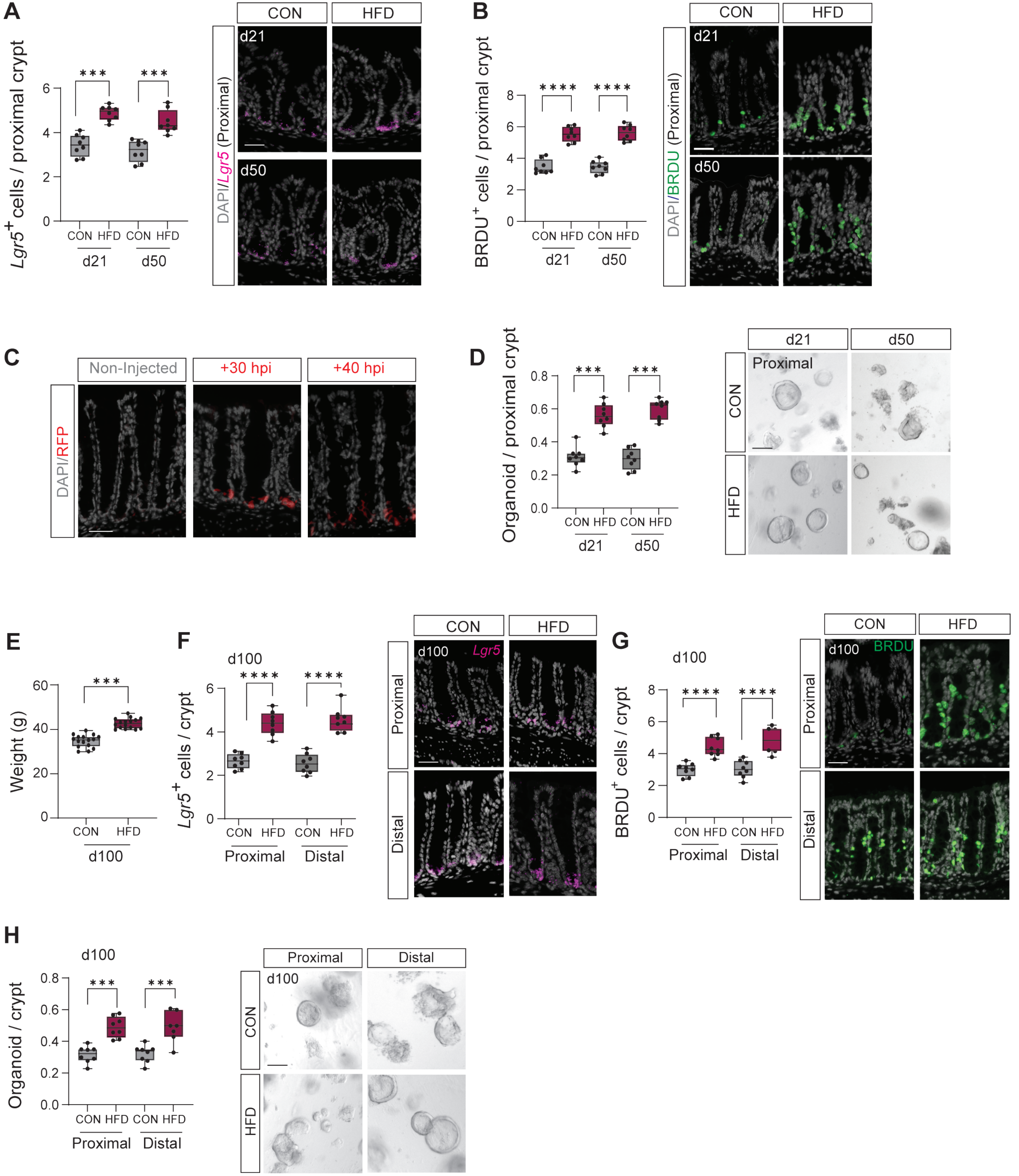

### Supplementary Figure 2

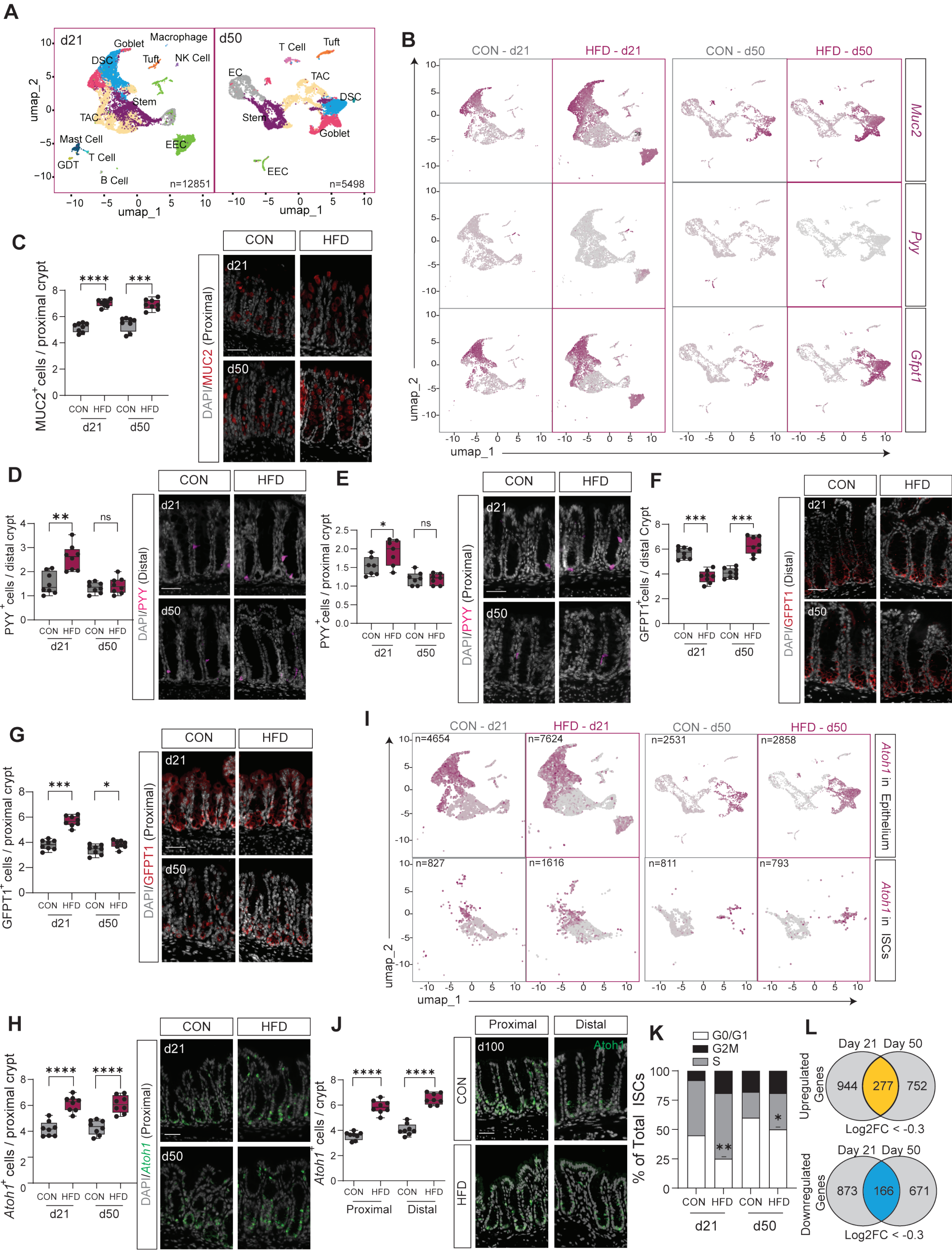

### Supplementary Figure 3

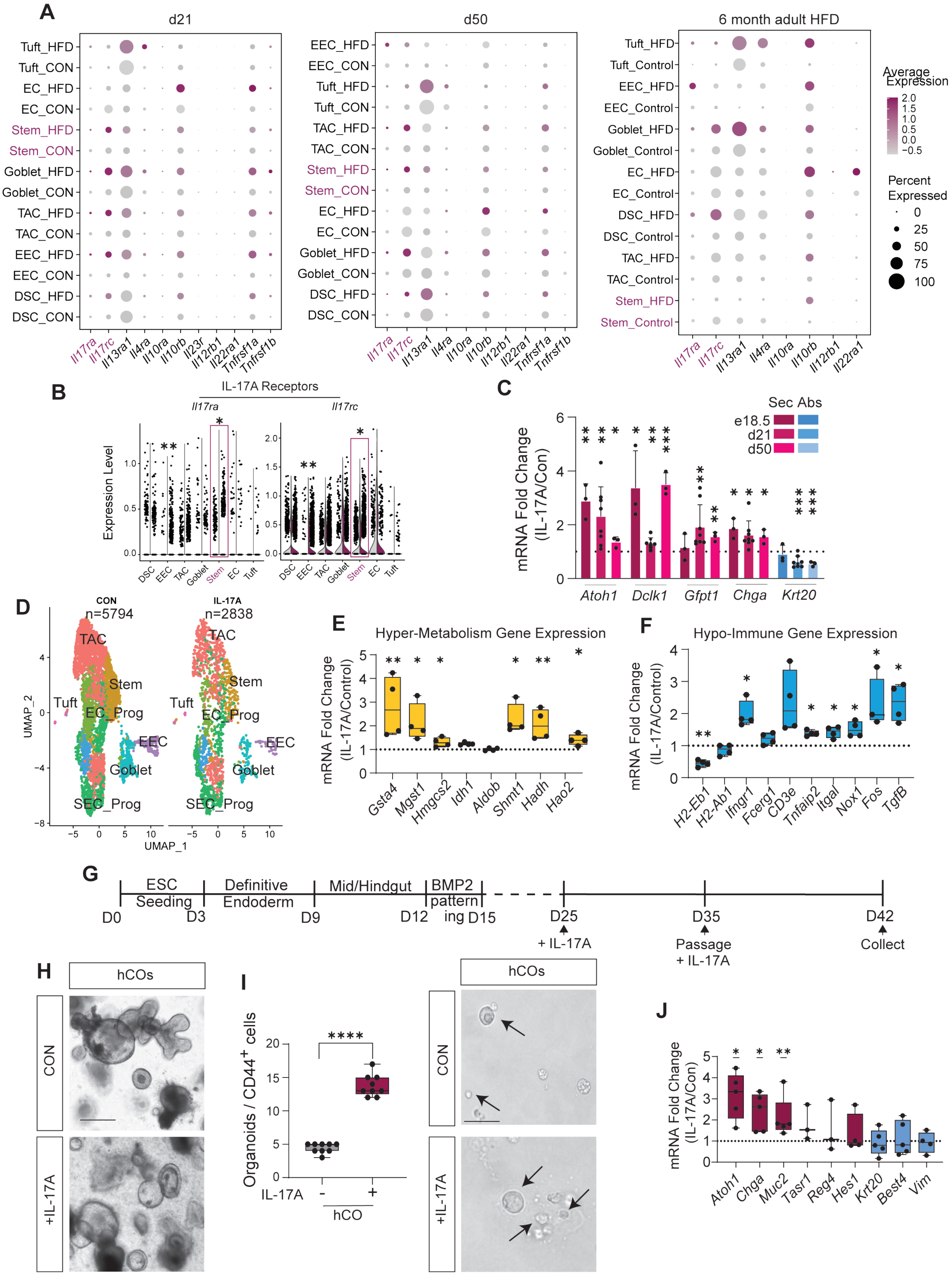

### Supplementary Figure 5

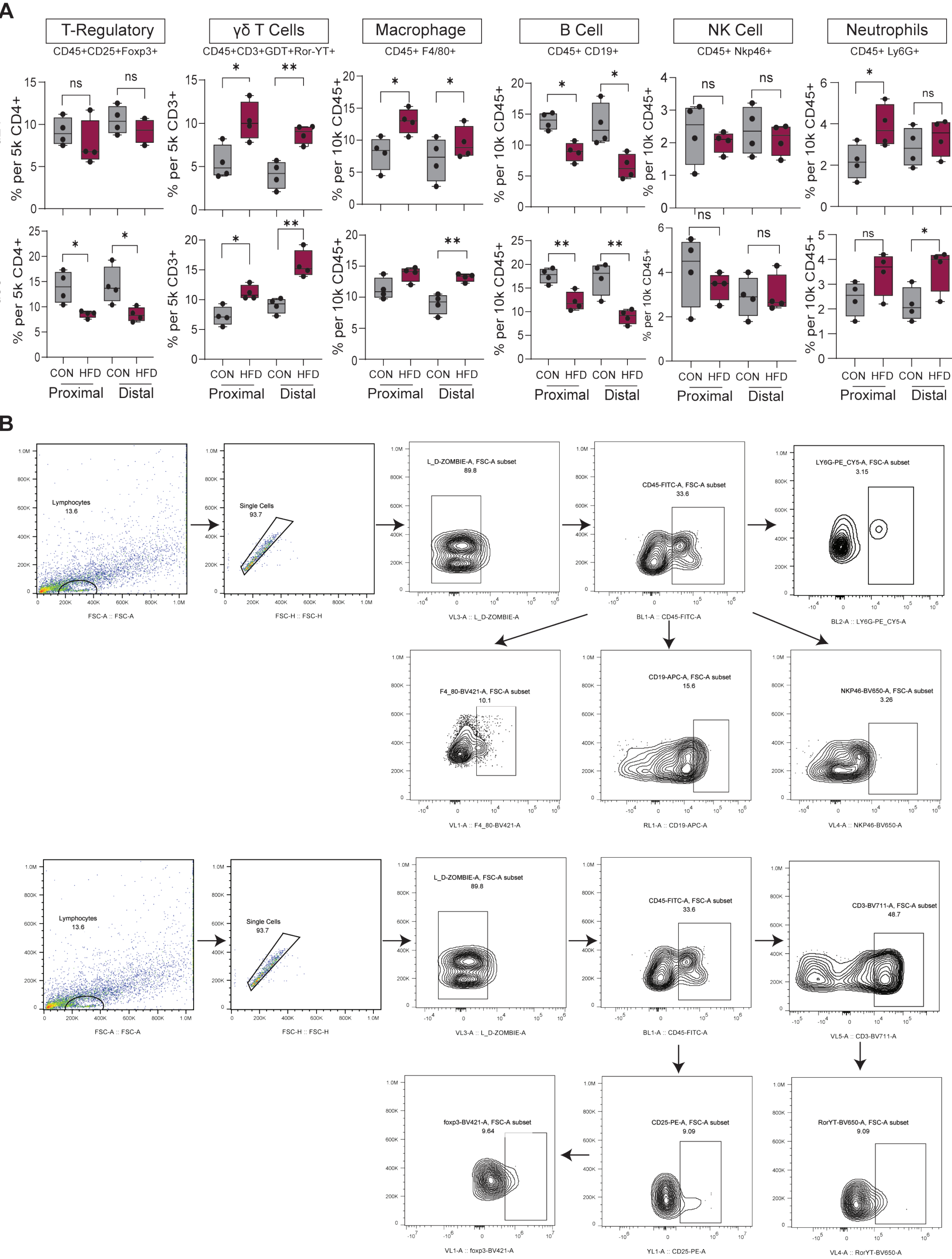

### Supplementary Figure 6

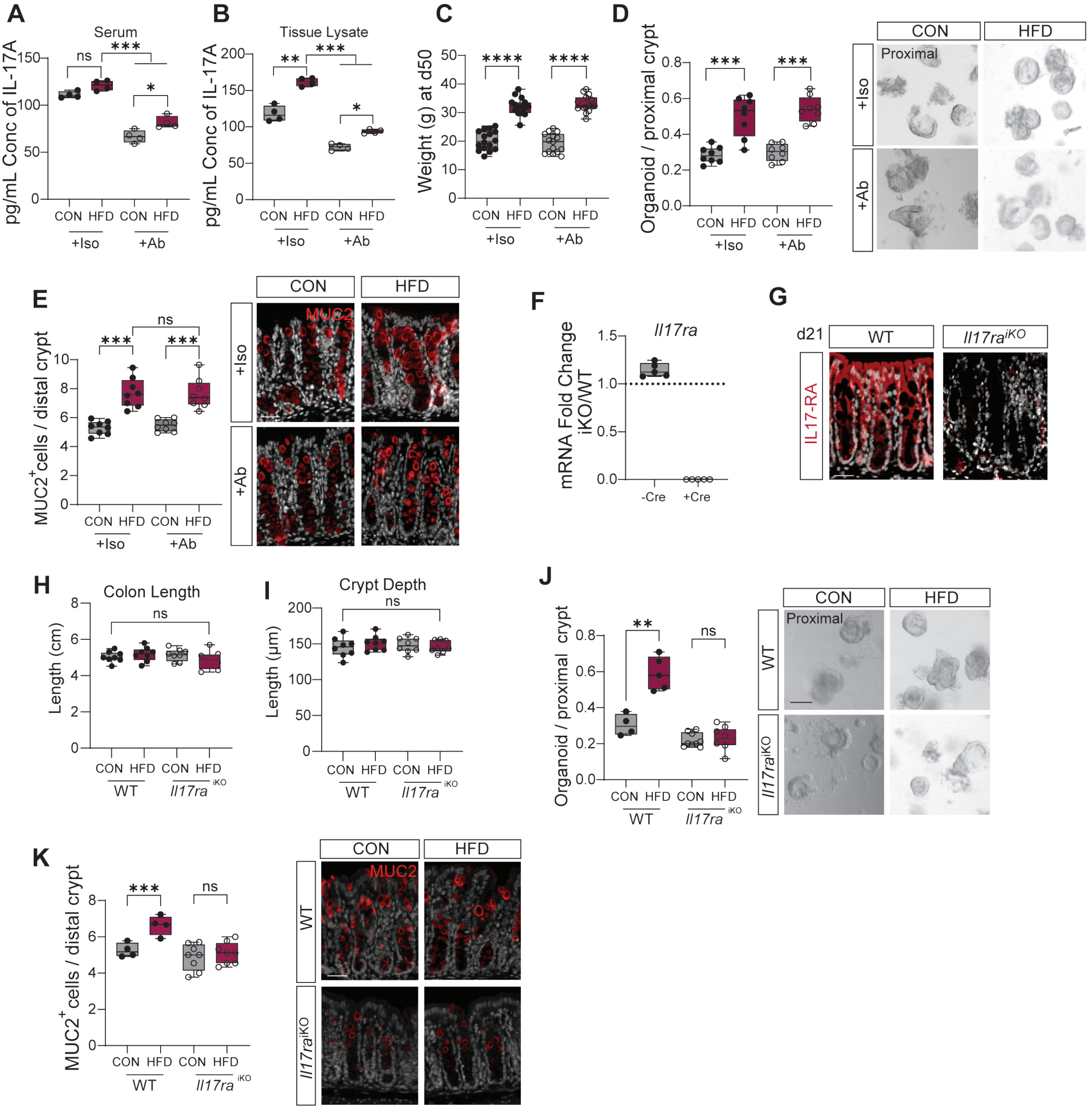

### Supplementary Figure 7

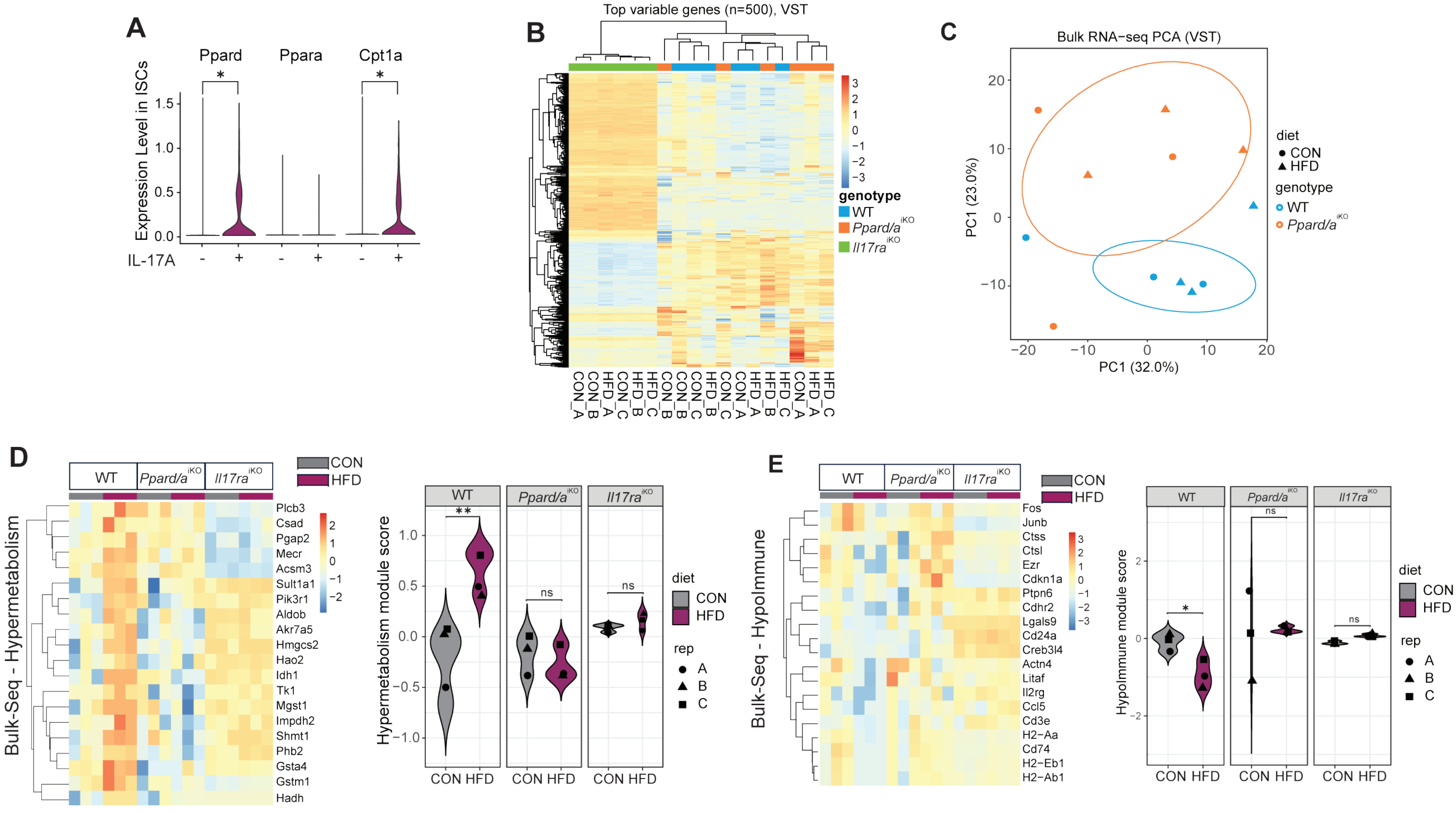
