## Supplementary Figure 4 for "Maternal obesity induces developmental programming of Intestinal stem cells through an IL-17A/PPAR immune-epithelial axis"

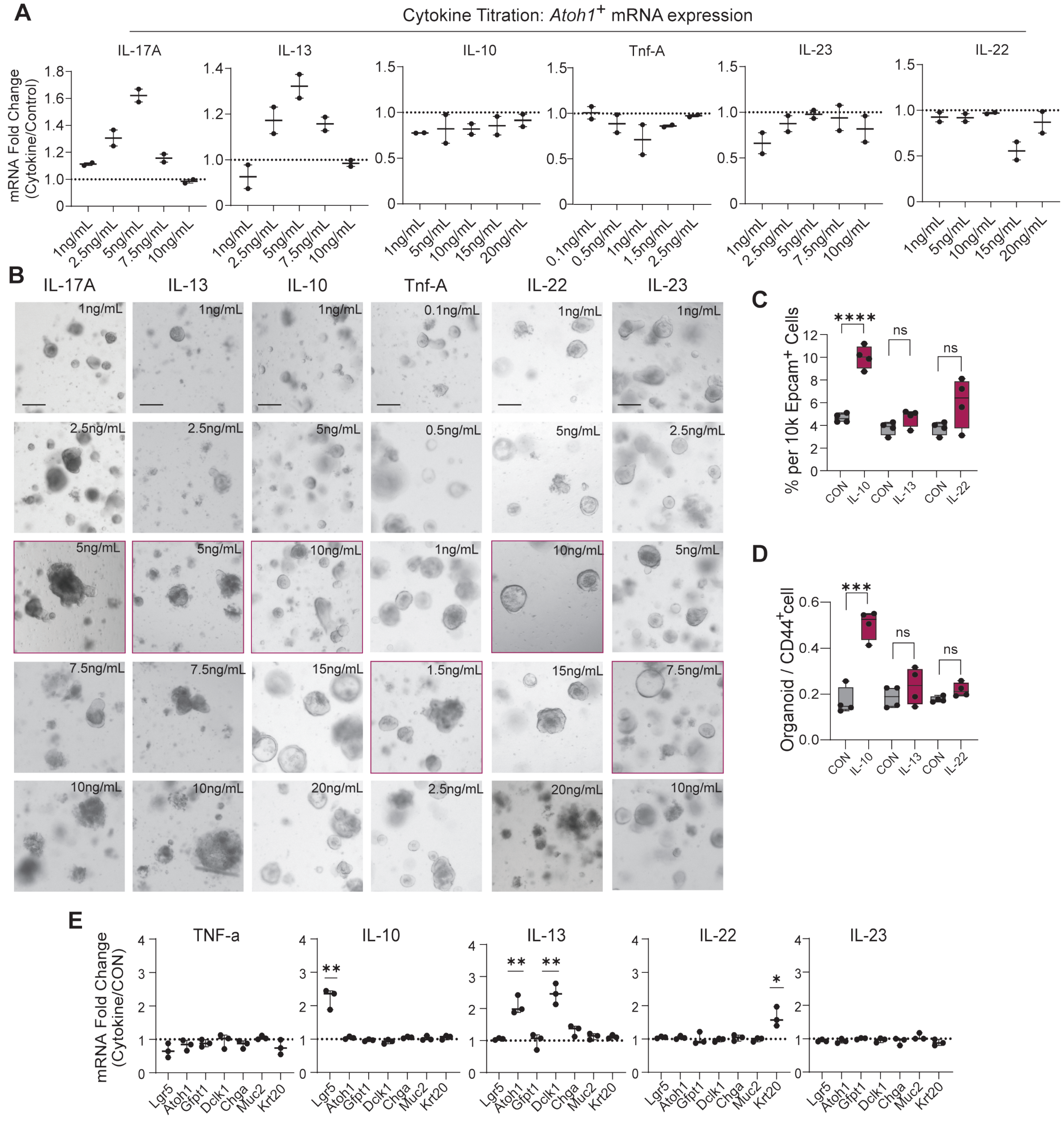


**F**

| Cytokine | Concentration | Stem Cell CD44+ Freq | Clonogenicity | Atoh1 Expression | Secretory Lineage | Rationale |
|---|---|---|---|---|---|---|
| IL-17A | 5ng/mL | Increase | Increase | Increase | Increase | Receptor freq increases in ISCs, Progs and Sec cells |
| IL-13 | 5ng/mL | No Change | No Change | Increase | Increase | Receptor freq increases in ISCs and Tufts cells |
| IL-22 | 10ng/mL | No Change | No Change | No Change | No Change | Receptor increases in ISCs |
| IL-10 | 10ng/mL | Increase | Increase | No Change | No Change | Major anti-inflammatory cytokine (negative control) |
| IL-23 | 7.5ng/mL | NA | NA | No Change | No Change | Pro-inflammatory cytokine functioning up of IL-17a. Target of multiple approved therapeutic agent |
| Tnf-A | 1.5ng/mL | NA | NA | No Change | No Change | Pro-inflammatory cytokine often works with IL-17a. Target of multiple approved therapeutic agent |
