## Supplementary Figure Legends for "Maternal obesity induces developmental programming of Intestinal stem cells through an IL-17A/PPAR immune-epithelial axis"

**Supplemental Figure 1: Maternal obesogenic conditions induce persistent effects in offspring ISCs**
(A) Quantification and representative images of *Lgr5*$^{+}$ ISC smFISH per crypt from the proximal colon at d21 and d50. Each point represents the mean of >20 crypts analyzed per mouse (n=8). Scale bar = 20μm
(B) Quantification and representative images of BRDU$^{+}$ crypt cells from the proximal colon at d21 and d50. Each point represents the mean of >20 crypts analyzed per mouse (n=8). Scale bar = 20μm
(C) Representative images of Tamoxifen IP injections post 30 and 40 hours. hpi: hours post injection.
(D) Clonogenicity of proximal crypts from CON and HFD d21 mice. Each point represents the mean of 3+ wells from an individual animal. Scale bar = 10μm.
(E) Animal weight at d100 (n=16 each).
(F) Quantification and representative images of *Lgr5*$^{+}$ ISC smFISH per crypt from the proximal colon at d100. Each point represents the mean of >20 crypts analyzed per mouse (n=8). Scale bar = 20μm
(G) Quantification and representative images of BRDU$^{+}$ cells crypt from the proximal colon at d100. Each point represents the mean of >20 crypts analyzed per mouse (n=8). Scale bar = 20μm
(H) Clonogenicity of proximal and distal crypts from CON and HFD-exposed mice at d100. Each point represents the mean of 3+ wells from an individual animal. Scale bar = 10μm.

**Supplemental Figure 2: mHFD exposure shifts lineage bias towards a secretory phenotype**
(A) UMAP of the epithelial and immune cell clusters in offspring at d21 and d50.
(B) UMAP of *Muc2*, *Pyy* and *Gfpt1* expression of in offspring at d21 and d50.
(C) Quantification and representative images of MUC2$^{+}$ goblet cells per proximal colonic crypt at d21 and d50. Each point represents the average mean of >20 crypts analyzed per mouse (n=8). Scale bar = 20μm
(D-G) Quantification and representative images of *Pyy*$^{+}$ enteroendocrine cells and *Gfpt1*$^{+}$ deep secretory cells by IF per (D,F) distal and (E,G) proximal colonic crypt at d21and d50. Each point represents the average mean of >20 crypts analyzed per mouse (n=8). Scale bar = 20μm
(H) Quantification and representative images of *Atoh1*$^{+}$ transcripts per proximal colonic crypt at (I) d21 and d50. Each point represents the mean of >20 crypts analyzed per mouse (n=8). Scale bar = 20μm
(I) Feature plot showing the *Atoh1*$^{+}$ cells in ISCs at d21 and d50.
(J) Quantification and representative images of *Atoh1*$^{+}$ transcripts per proximal colonic crypt at d100. Each point represents the mean of >20 crypts analyzed per mouse (n=8). Scale bar = 20μm
(K) Stacked bar plot of ISCs cell-cycle analysis using Seurat's CellCycle at d21 and d50.
(L) Venn diagram of the commonly upregulated and downregulated genes in ISCs at d21 and d50.

**Supplemental Figure 3: IL-17A cytokine promotes mHFD phenotypes**
(A) Dotplot of cytokine receptor expression profiles in the distal colon of d21 and d50 offspring from maternal dietary intervention and whole colon of adult-induced 6-month HFD
(B) Violin plot of *Il17ra* and *Il17rc* expression across distal colonic epithelial cell populations at d21 and d50
(C) qRT-PCR analysis of epithelial cell population from Control and IL-17A-treated distal colon organoids derived from e18.5, d21 and d50 mice. Each point represents the normalized CT value (IL-17A/CON) per mouse (e18.5 n=3, d21 n=8, d50, n=3).
(D) UMAP of Control and IL-17A-treated e18.5 colon organoids.
(E, F) qRT-PCR analysis of Hypermetabolism (D) and Hypoimmune (E) genes from Control and IL-17A treated distal colon organoids derived from d21 WT mice. Each point represents the normalized CT value (IL17A/CON) per mouse (n=4).
(G) Schematic depicting the generation and treatment of hESC derived colon organoids.
(H) Representative images of Control and IL-17A-treated hCOs at d42.
(I) Clonogenicity of CD44$^{+}$ sorted cells from hCOs with representative images, day6 post plating. Each point represents the mean of 3+ wells per hCO. Scale bar = 10μm
(J) qRT-PCR analysis of Control and IL-17A-treated hCOs. Each point represents the normalized CT value (IL-17A/CON) per hCO (n=5).

**Supplemental Figure 4: *In vitro* cytokine treatment on WT distal colon organoids**
(A) Cytokine titration and qRT-PCR analysis of *Atoh1*$^{+}$ expression in mouse d21 distal colon organoids (n=2).

(B) Brightfield organoid images of distal colon organoids post cytokine treatment. Images were taken on d9 post plating of crypts (see schematic Figure 3E).
(C) Quantification by flow cytometry of $CD44^+$ cells from Control and cytokine-treated distal colon organoids (n=4).
(D) Clonogenicity of $CD44^+$ sorted cells post cytokine treatment. Each point represents the mean of 3+ wells per mouse and per cytokine.
(E) qRT-PCR analysis of Control and cytokine-treated mouse distal colon organoids derived from d21 mouse. Each point represents the normalized CT value (Cytokine/CON) per mouse (n=3).
(F) Summary table showing the outcome of cytokine treatments on mouse distal colon organoids.

**Supplemental Fgure 5: mHFD exposure alters the colonic immune landscape**
(A) Quantification of colonic immune cells at d21 and d50 across maternal dietary conditions (n=4).
(B) Flow density plot showing the gating strategy to identify the immune populations.

**Supplemental Figure 6: *IL17ra*-deficiency blocks the mHFD phenotype**
(A) Serum IL-17A levels at d50 via ELISA post Isotype/Anti-IL-17A treatment (n=4).
(B) Tissue lysate IL-17A levels at d50 via ELISA post Isotype/Anti-IL-17A treatment (n=4).
(C) Weight from Isotype and Antibody treated offspring at d50 (n=15).
(D) Clonogenicity of proximal colonic crypts with representative images. Each point represents the mean of 3+ wells per mouse. Scale bar = 10µm
(E) Quantification and representative images of $MUC2^+$ goblet cells per distal colonic crypt. Each point represents the mean of >20 crypts analyzed per mouse (n=8). Scale bar = 20µm
(F, G) qRT-PCR (F) and IF validation (G) of IL-17RA depletion in *Il17ra*$^{iKO}$ offspring compared to WT controls.
(H, I) Quantification of colon length (H) and crypt depth (I) in WT and *Il17ra*$^{iKO}$ offspring at d21 (n=8).
(J) Clonogenicity of proximal colonic crypts with representative images from crypts in WT and *Il17ra*$^{iKO}$ offspring at d21. Each point represents the mean of 3+ wells per mouse. Scale bar = 10µm
(K) Quantification and representative images of $MUC2^+$ goblet cells from the distal colon in WT and *Il17ra*$^{iKO}$ offspring at d21. Each point represents the mean of 3+ wells per mouse. Scale bar = 10µm

**Supplemental Figure 7: An IL-17A-PPARd/a axis patterns early ISCs**
(A) Violin plot showing the expression of *Ppar-d*, *Ppar-a* and *Cpt1a* in ISCs from e18.5 organoids treated with Control and IL-17A.
(B) Heatmap from BulkSeq analysis showing the z-scored expression of top 500 variable gene expression profile from d21 mCON and mHFD offspring crypts of WT, *Ppar-d/a*$^{iKO}$ and *Il17ra*$^{iKO}$
(C) PCA plot of variance-stabilized gene expression profiles of crypts derived from d21 WT and *Ppar-d/a*$^{iKO}$ and offspring.
(D,E) Heatmap analysis showing z-scored hypermetabolic (D) and hypoimmune (E) gene expression profiles from bulk RNA-Seq of d21 mCON and mHFD crypts derived from WT, *Ppar-d/a*$^{iKO}$ and *Il17ra*$^{iKO}$ animals.
